# Phenotype-specific melanoma uptake of fatty acid from human adipocytes activates AXL and CAV1-dependent β-catenin nuclear accumulation

**DOI:** 10.1101/2024.01.21.576568

**Authors:** Ana Chocarro-Calvo, Miguel Jociles-Ortega, José Manuel García-Martinez, Pakavarin Louphrasitthiphol, Yurena Vivas Garcia, Ana Ramírez-Sánchez, Jagat Chauhan, M Carmen Fiuza, Manuel Duran, Custodia García-Jiménez, Colin R. Goding

**Author notes:** Corresponding Authors: Colin R Goding, Custodia García Jiménez, Te1. Equal contribution (Ana Chocarro-Calvo, Miguel Jociles-Ortega).

## Abstract

Phenotypic diversity of cancer cells within tumors generated through bi-directional interactions with the tumor microenvironment has emerged as a major driver of disease progression and therapy resistance. Nutrient availability plays a critical role in determining phenotype, but whether specific nutrients elicit different responses on distinct phenotypes is poorly understood. Here we show, using melanoma as a model, that only MITF^Low^ undifferentiated cells, but not MITF^High^ cells, are competent to drive lipolysis in human adipocytes. In contrast to MITF^High^ melanomas, adipocyte-derived free fatty acids are taken up by undifferentiated MITF^Low^ cells via a fatty acid transporter (FATP)-independent mechanism. Importantly, oleic acid (OA), a monounsaturated long chain fatty acid abundant in adipose tissue and lymph, reprograms MITF^Low^ undifferentiated melanoma cells to a highly invasive state by ligand-independent activation of AXL, a receptor tyrosine kinase associated with therapy resistance in a wide range of cancers. AXL activation by OA then drives SRC-dependent formation and nuclear translocation of a β-catenin-CAV1 complex. The results highlight how a specific nutritional input drives phenotype-specific activation of a pro-metastasis program with implications for FATP-targeted therapies.

## Introduction

Tumors contain multiple, phenotypically distinct cell subpopulations that share common driver mutations but exhibit radically different biological properties. Specifically, in response to changing microenvironmental cues, phenotypically plastic cells may reversibly switch *in vivo* from one phenotype to another (Hanahan, 2022; Hoek and Goding, 2010). The ability of cells to change phenotype represents a major challenge to effective anti-cancer therapy by underpinning metastatic dissemination and contributing to both drug- and immunotherapy-tolerance (Gerstberger et al., 2023; Kalkavan et al., 2022; Sharma et al., 2010). Although there have been major advances in our understanding of how microenvironmental cues impose specific phenotype switches, much less is known about whether or how specific phenotypic states exhibit distinct responses to the same microenvironment. Understanding phenotype-specific responses will be crucial to the development of novel and more effective anti-cancer therapies. (Rambow et al., 2019).

Although the intratumor microenvironment is complex, one key determinant of cell behaviour is nutrient availability (García-Jiménez and Goding, 2019). To facilitate tumor expansion, cancers induce growth of new blood vessels to increase nutrient and oxygen supply (De Palma et al., 2017). However, the chaotic vasculature within tumors means that oxygen and nutrients are frequently limiting. Since proliferation is critically dependent on nutrient availability, instructive cues that trigger cell division also coordinate nutrient uptake and processing (Pavlova and Thompson, 2016). In cancer, activated oncogenes may bypass the need for pro-proliferative signals and re-wire cellular metabolism (Pavlova and Thompson, 2016). However, without a sufficient supply of nutrients, oncogenes cannot drive cell division. To compensate for limited nutrient availability, cells restrict nutrient demand, for example by reducing protein translation and slowing the cell cycle (Falletta et al., 2017; Lee et al., 2021; Vivas-Garcia et al., 2020), and implement strategies that increase nutrient supply(García-Jiménez and Goding, 2019). These include adopting an invasive phenotype so as to search for new nutrient supplies, promoting new vessel formation, increasing uptake of nutrients and engaging autophagy to recycle damaged organelles (García-Jiménez and Goding, 2019).

An early response of cells lacking key resources is to trigger metabolic symbiosis by promoting nutrient release from neighboring cells (Lyssiotis and Kimmelman, 2017; Martinez-Outschoorn et al., 2014; Nakajima and Van Houten, 2013; Sonveaux et al., 2008; Sousa et al., 2016). Adipocytes are designed physiologically to store excess energy as lipids and release it when required. Increasing evidence suggests this process is subverted in cancer, and tumor-associated adipocytes can release lipids in response to signals from tumor cells (Hoy et al., 2021). Lipids released by adipocytes can be used to obtain energy or as a carbon source to fuel anabolism in proliferating cancer cells (Hoy *et al*., 2021). In this respect, invasive cancer cells have frequently been observed in close proximity to adipocytes (Muller et al., 2013). For example, in ovarian cancer lipid transfer from adipocytes in the fat-rich omentum increases cancer cell proliferation (Nieman et al., 2011), while in breast cancer, lipid release from adipocytes promotes not only cancer cell proliferation (Balaban et al., 2017) but also invasion (Dirat et al., 2011).

Melanoma is a highly aggressive skin cancer with well-defined phenotypic subtypes (Goodall et al., 2008; Hoek and Goding, 2010; Rambow *et al*., 2019; Rambow et al., 2018; Tsoi et al., 2018), initially distinguished by the expression of the Microphthalmia-associated transcription factor MITF (Goding and Arnheiter, 2019) that controls many aspects of melanoma biology including phenotypic identity. Melanoma therefore represents an excellent model to explore how lipid uptake has consequences for phenotypic identity and disease progression. Cutaneous melanoma usually originates in the basal layer of the epidermis, and as tumors develop melanoma cells can come into contact with the adipocyte-rich dermis (Kwan et al., 2014). Consequently, adipocyte association with tumors is a marker of poor prognosis (Smolle et al., 1995), with adipocyte-derived exosomes promoting fatty acid oxidation and invasion in melanoma cells (Lazar et al., 2016). Importantly, in xenograft melanoma models peri-tumoral adipocytes exhibit reduced size and characteristics of increased lipolysis (Wagner et al., 2012). These observations are consistent with melanomas inducing lipolysis in adipocytes in culture (Hollander et al., 1986; Kwan *et al*., 2014). Moreover, recent compelling evidence from both in vitro and in vivo models indicate that fatty acids released from stromal adipocytes are imported by melanoma cells using fatty acid transporter proteins (FATPs) to fuel disease progression (Lumaquin-Yin et al., 2023; Zhang et al., 2018). Significantly, oleic acid uptake from lymph can protect metastasizing melanoma cells from ferroptosis (Ubellacker et al., 2020) and FATP-dependent lipid uptake from aged fibroblasts can contribute to tolerance to BRAF inhibitor therapy (Alicea et al., 2020). However, while it is clear that fatty acid uptake plays a key role in melanoma biology, mechanistically how it rewires melanoma signaling to achieve its biological effects are poorly understood. Neither is it understood whether melanoma phenotype dictates the molecular response to specific fatty acids nor whether the release of fatty acids from human adipocytes is a response to signals restricted to specific phenotypes.

Here we reveal that only MITF^Low^ melanoma cells induce lipolysis in human adipose tissue explants, with adipocyte lipolysis being triggered by melanoma-derived WNT5a. Surprisingly, we also find that FATP-dependent uptake of fatty acids is restricted to MITF^High^ melanoma cells. In contrast, undifferentiated MITF^Low^ AXL^High^ therapy-resistant melanoma cells use a FATP-independent but AXL-dependent mechanism to take up fatty acids, leading to AXL-dependent activation of SRC, phosphorylation of Caveolin 1, translocation of a Caveolin-β-catenin complex to the nucleus and increased invasion.

## Results

### Lipolysis of human adipose tissue explants is induced only by MITF^Low^ melanoma

In vivo, metastasizing cancer cells can come into close proximity with adipocytes with the potential for bidirectional interactions including cancer cell-induced adipocyte lipolysis and uptake of the released fatty acids by the cancer cells. However, whether human adipocytes release fatty acids in response to signals originating from specific phenotypic states is unknown. To investigate this, we used two BRAF mutant human melanoma cell lines, IGR37 and IGR39 that are derived from the same patient but which are phenotypically very different: IGR37 cells express MITF and represent a proliferative phenotype that is BRAF inhibitor (BRAFi) sensitive; by contrast IGR39 cells are MITF^Low^, undifferentiated, invasive, and BRAFi-tolerant, and express the AXL receptor tyrosine kinase, a hallmark of therapy resistance. The interaction between melanoma cells and adipocytes was recapitulated by co-culture with human adipose tissue explants separated by a membrane that permits lipid exchange but does not allow passage of cells (Figure 1A). Fatty acid uptake by the melanoma cells was then monitored using Nile red, a fluorescent lipophilic dye. Unexpectedly, only the undifferentiated MITF^Low^ IGR39 cells, and not the MITF^High^ IGR37 cells accumulated Nile red-stained lipid droplets (Figure 1B). This result might be obtained because only the IGR39 cells were competent to induce lipolysis in the human adipose tissue explants, and/or because only IGR39 cells could take up the lipids released from the explants.

**Figure 1.**
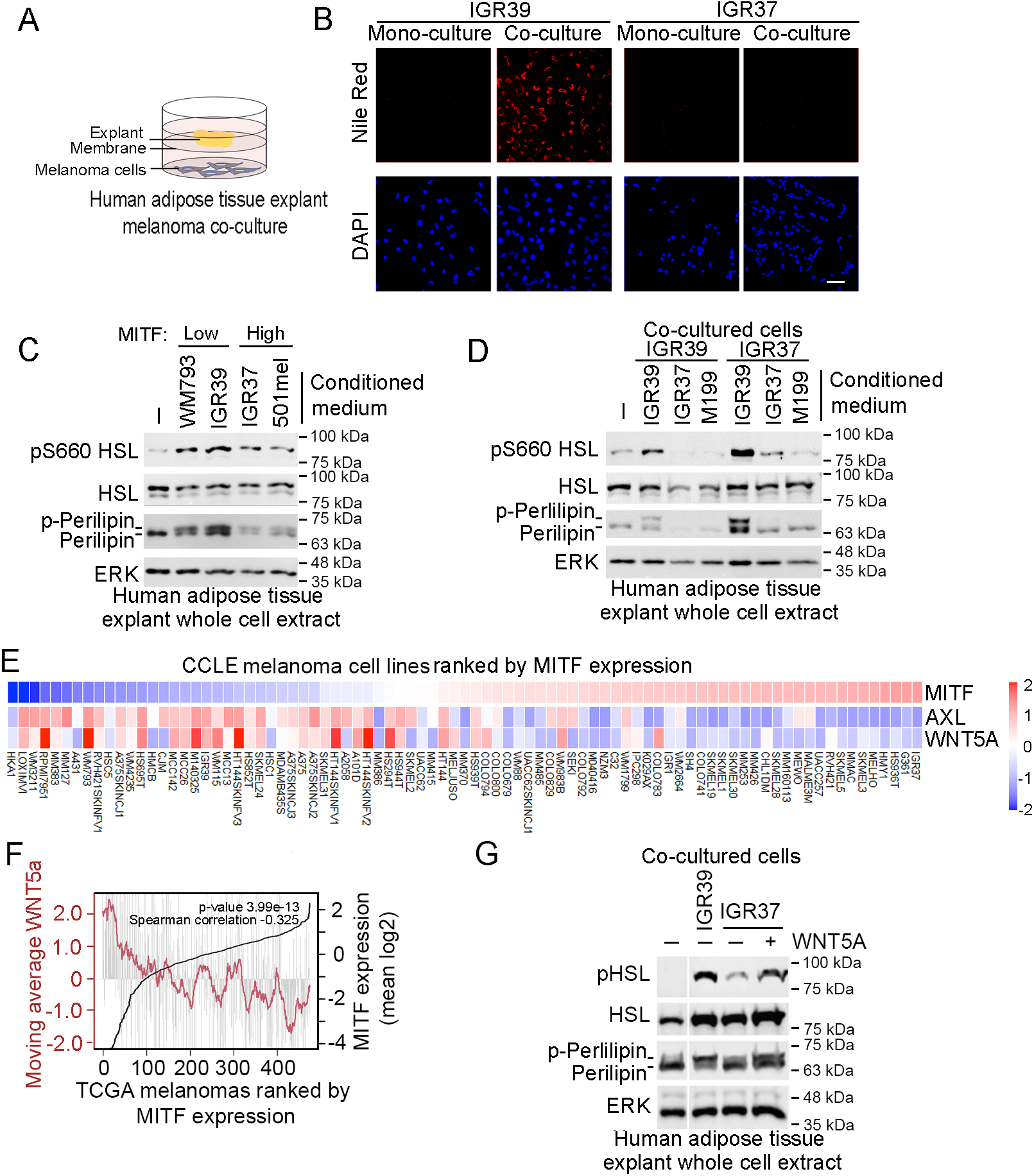
Only MITF^Low^ cells induce lipolysis in human adipose explants. **A.** Schematic showing experimental system for adipose tissue explant-melanoma cell co-culture in which the explant and melanoma cells are separated by a lipid permeable membrane. **B.** Nile red staining of melanoma cells in mono-culture or co-culture with human adipose tissue explants for 4 days. DAPI is used to stain nuclei. Scale bar = 50 μM. **C, D.** Western blots using indicated antibodies of human adipose tissue explants co-cultured with indicated melanoma cells and exposed to indicated conditioned medium for 4 days. **E.** Heatmap showing relative mRNA expression of indicated genes in CCLE melanoma cell lines ranked by *MITF* expression. **F**. Plot showing *WNT5a* mRNA expression (grey bars) in the TCGA melanoma cohort ranked by *MITF* expression (black line). The moving average per 20 melanomas of *WNT5a* is shown as the maroon line. **G**. Western blot of a human adipose tissue explants co-cultured or not with indicated cell lines +/-50 ng WNT5a. All samples were run on the same gel.

To distinguish between these possibilities, we initially examined the ability of both IGR39 and IGR37 cells to induce phosphorylation of the hormone-stimulated lipase (HSL), a key activation event in mobilization of adipose tissue fat stores. To ensure that the results were not limited to IGR37 and IGR39 cells, we used two additional melanoma cell lines: WM793, that like IGR39 are MITF^Low^, undifferentiated cells; and 501mel cells, that like IGR37 exhibit an MITF^High^, proliferative phenotype. Human adipose tissue explants exposed to conditioned medium derived from the 4 melanoma cell lines for 4 days were examined for HSL phosphorylation status by Western blot. The results (Figure 1C) revealed that conditioned medium from both undifferentiated melanoma cell lines (WM793 and IGR39) induced phosphorylation of HSL, but not that from the MITF^High^ IGR37 and 501mel cells. In addition, conditioned medium from WM793 and IGR39 cells, but not that from IGR37 or 501mel cells induced expression and phosphorylation of perilipin, a lipid droplet-associated protein and gatekeeper of lipolysis, whose phosphorylation is a hallmark of adipocyte lipolysis. ERK was used as a loading control.

To extend these observations, we co-cultured IGR37 or IGR39 cells with human adipose tissue explants in conditioned medium from IGR39 cells, IGR37 cells or in M199 adipocyte culture medium. The results confirmed that co-cultured undifferentiated IGR39 cells could induce both HSL and perilipin phosphorylation, which was reduced in the presence of IGR37 conditioned medium or M199 medium. By contrast cocultured IGR37 cells could not induce markers of lipolysis in the human adipose tissue explants unless cells were exposed to conditioned medium from IGR39 cells. Collectively, these results indicate that only factors secreted from undifferentiated melanoma cells could induce lipolysis in human adipose tissue explants.

To rule out any defects in uptake of free fatty acids by IGR37 cells, we next exposed IGR37 and IGR39 cells to oleic acid, an 18:1 long chain monounsaturated fatty acid, and examined the accumulation of lipid droplets. Oleic acid was used since it is the most abundant fatty acid in human adipose tissue (Hodson et al., 2008; Insull and Bartsch, 1967; Kokatnur et al., 1979) and previous work has shown that uptake of oleic acid from lymph can enhance survival and suppress ferroptosis in metastasizing melanoma cells (Ubellacker *et al*., 2020). In these experiments we used 100 μM oleic acid, a concentration 5-fold lower than that used previously to suppress ferroptosis (Ubellacker *et al*., 2020). As expected, exposure of both cell lines to oleic acid led to accumulation of lipid droplets revealed by staining with Oil Red (Supplementary Figure 1A).

These observations indicate that while IGR37 cells cannot induce lipolysis in human adipose tissue explants, they are nevertheless capable of importing free fatty acids. We therefore co-cultured IGR37 and IGR39 cells separated by a porous membrane with murine 3T3-L1 adipocytes preloaded with fluorescent BODIPY-FL that acts as a fatty acid mimetic and which is transported by the same mechanisms as natural lipid (Bai and Pagano, 1997) (Supplementary Figure 1B). Any uptake of BODIPY-FL by co-cultured melanoma cells therefore reflects both its release from the adipocytes as well as an ability of the melanoma cells to import it. Unlike the result obtained with human adipose tissue explants, IGR37 and IGR39 cells were both able to induce BODIPY-FL release from the 3T3-L1 cells and its uptake (Supplementary Figure 1C). Taken together our results indicate that both MITF^Low^ and MITF^High^ cells can take up oleic acid, but that while both can induce lipid release and uptake from the murine 3T3-L1 adipocyte, only undifferentiated MITF^Low^ cells are competent to induce lipolysis from human adipose tissue explants and import the lipids released. Collectively, these data suggest that a soluble factor secreted from MITF^Low^, but not MITF^High^, melanoma cells is able to induce lipolysis and lipid release from human adipose tissue explants. Although a range of factors might be responsible, to date, no melanoma secreted factor able to induce lipid release from adipocytes has been identified. However, a number of previous studies suggest that WNT5a, an activator of the of non-canonical WNT signalling pathway and driver of melanoma invasion (Weeraratna et al., 2002) and BRAF inhibitor resistance (Anastas et al., 2014), might be a potential candidate. First, circulating WNT5a can promote a pro-inflammatory state in visceral adipose tissue (Catalan et al., 2014) and can drive adipocyte dedifferentiation associated with loss of lipid droplets (Zoico et al., 2016). Second, analysis of the panel of melanoma cell lines in the Cancer Cell Line Encyclopedia (CCLE) revealed that WNT5a was primarily expressed in MITF^Low^ cell lines (including IGR39 and WM793 cells) (Figure 1E), using the expression of the AXL receptor tyrosine kinase (RTK) as an additional marker of MITF^Low^, therapy resistant phenotype (Muller et al., 2014). The results obtained in cell lines were recapitulated in the TCGA melanoma cohort in which *WNT5a* was expressed predominantly in MITF^Low^ tumors (Figure 1F), consistent with both low MITF (Carreira et al., 2006; Chauhan et al., 2022) and high WNT5a (Weeraratna *et al*., 2002) being associated with invasiveness. We therefore tested the ability of WNT5a to induce lipolysis in human adipose tissue explants. The results (Figure 1G) confirmed that both p-Perilipin and p-HSL were robustly induced by co-culture with IGR39 cells. Although co-culture with IGR37 cells rendered a moderate increase in lipolysis, addition of WNT5a to the IGR37 cells led to a substantial increase. These data are consistent with WNT5a secreted by MITF^Low^, but not MITF^High^, cells contributing to their ability to induce lipolysis in human adipose tissue.

### Uptake of fatty acids by MITF^Low^ melanomas is FATP-independent

The observation that only undifferentiated melanoma cells were competent to induce effective lipolysis from human adipose tissue explants led us to reassess how fatty acids are transferred into melanoma cells. Previous work has demonstrated that fatty acid uptake into melanoma cells from adipocytes or aged fibroblasts is mediated by FATPs and can contribute to therapy resistance (Alicea *et al*., 2020; Zhang *et al*., 2018). However, whether FATPs are also required for fatty acid uptake by undifferentiated melanoma cell lines has not been examined. We therefore exposed two MITF^High^ (501mel and IGR37) and two MITF^Low^ (IGR39 and WM793) human melanoma cell lines to oleic acid in the presence or absence of Lipofermata, a small molecule that specifically blocks the ability of FATPs to import long-chain fatty acids (Sandoval et al., 2010). The accumulation of intracellular lipid droplets within the melanoma cells was monitored using Nile red (Figure 2A; quantified in 2B). In the absence of exogenously added oleic acid, some melanoma cells may contain a few small lipid droplets likely originating from fatty acids present in the fetal calf serum used in the culture medium. By contrast, addition of oleic acid led to a dramatic increase in lipid droplet formation in all cell lines, though the level of accumulation in the MITF^Low^ (IGR39 and WM793) cell lines was moderately reduced. As anticipated, lipofermata substantially reduced lipid droplet accumulation in the MITF^High^ (501mel and IGR37) cell lines. Unexpectedly, lipofermata exhibited no effect on lipid droplet accumulation in the MITF^Low^ IGR39 and WM793 cell lines. This result raised the possibility that MITF^Low^ undifferentiated melanoma cell lines might use an alternative pathway to take up long-chain fatty acids. We therefore examined the mRNA expression of a series of FATPs encoded by the *SLC27A* gene family in our in-house panel of 12 melanoma cell lines. We also included *CD36*, a fatty acid translocase that is expressed on metastasis-initiating cells characterized by increased fatty acid metabolism in a number of cancers (Pascual et al., 2016), and compared its expression to that of MITF, a marker for the more proliferative/differentiated cells. Remarkably, each cell line expressed a different pattern of FATP or CD36 mRNA expression (Figure 2C). For example, *CD36* was only significantly expressed in the MITF^Low^ WM115 line, while *SLC27A2*, encoding FATP2, previously implicated in lipid uptake from fibroblasts(Alicea *et al*., 2020), was only found in two MITF^Low^ cell lines, IGR39 and CHL. By contrast *SLC27A1* encoding FATP1, reported to mediate lipid uptake by melanoma cells from adipocytes in vivo (Zhang *et al*., 2018), was more widely expressed, though predominantly in MITF^High^ cell lines. Notably, one cell line (WM793) appeared to express little of any FATP or *CD36* mRNA, consistent with its lipofermata-independent lipid droplet accumulation.

**Figure 2.**
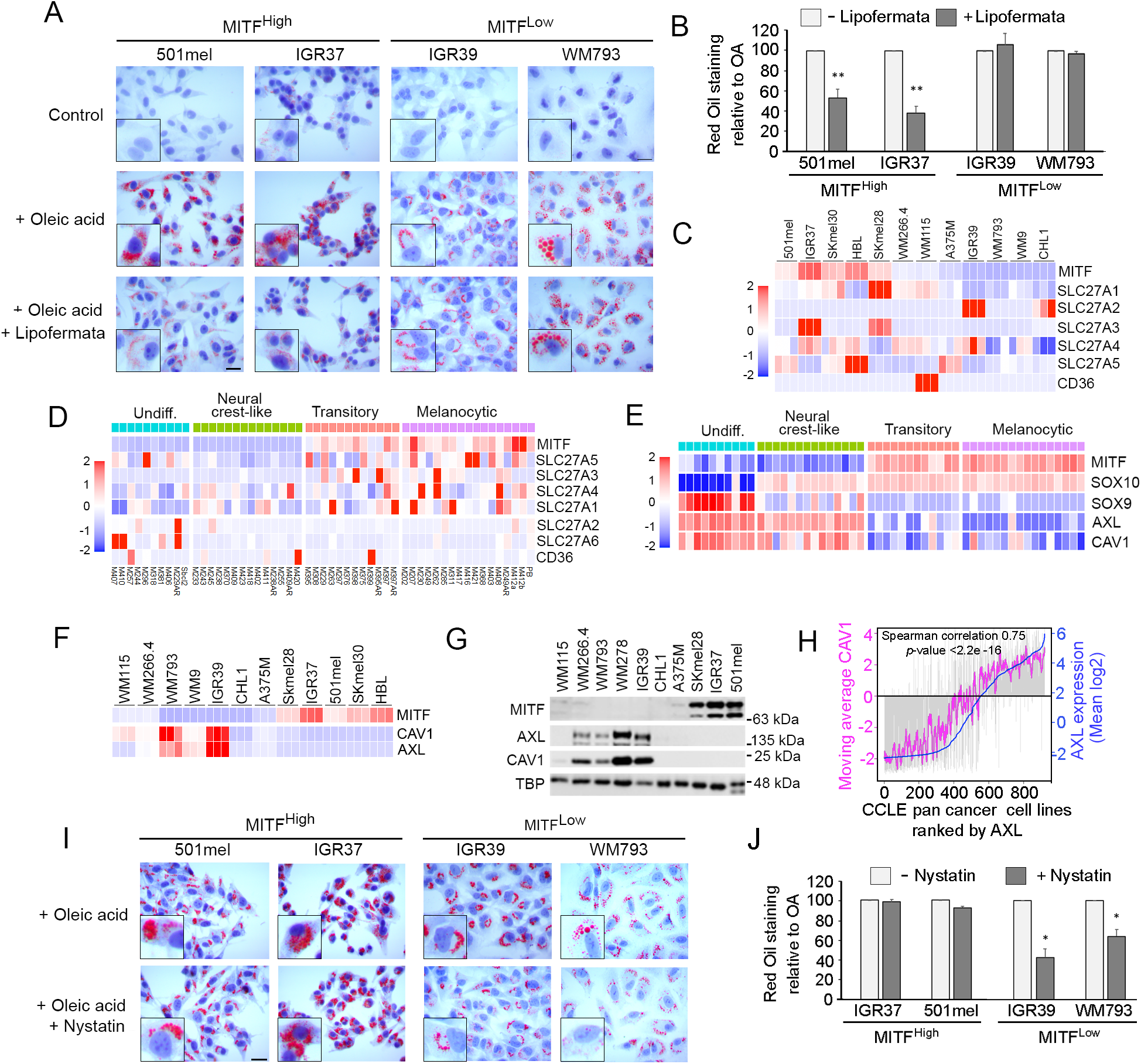
FATP-independent uptake of oleic acid by MITF^Low^ melanoma cells. **A.** Indicated MITF^High^ and MITF^Low^ cell lines exposed for 16 h or not to 100 μM oleic acid in the presence or absence of 1.25 μM Lipofermata and stained using Oil red O. Scale bars indicate 50 µm. Insets show typical cells at higher magnification. **B**. Quantification of Oil red O staining from cells in (A) n≥3, error bars indicate SEM * P<0.05. Paired t test. **C, D**. Heatmaps showing relative expression of MITF, CD36, and SLC27A family genes encoding FATPs in an in house panel of 12 melanoma cell lines measured by triplicate RNA-seq (C), or in the panel of 53 melanoma cell lines clustered by phenotype described in Tsoi et al. (2018) (D). **E**. Heatmap showing relative expression of *CAV1* in the Tsoi et al cell lines using *MITF*, *SOX10*, *SOX9* and *AXL* expression as markers of indicated phenotypes. **F**. Heatmap showing expression of *MITF*, *AXL* and *CAV1* mRNA in indicated panel of 12 melanoma cell lines by RNA-seq in triplicate. **G**. Western blot showing expression of indicated proteins in panel of 12 melanoma cell lines. **H.** CCLE pan cancer cell lines ranked by *AXL* expression with the moving average of *CAV1* expression indicated in magenta. Grey bars indicate *CAV1* expression in each cell line. **I**. Nile red staining of indicated melanoma cell lines exposed to oleic acid for 16 h, pre-treated or not with 25 μM Nystatin Insets indicate typical cells at higher magnification. Scale bar indicates 50 μm. **J**. Quantification of Nile red staining in (I) n≥3, error bars indicate SEM * P<0.05. Paired t test.

To extend these observations we next used a panel of 53 well characterized melanoma cell lines that fall into 4 distinct phenotypes based on their gene expression profiles (Tsoi *et al*., 2018). The results (Figure 2D) largely recapitulated those of our in-house cell lines, with the majority of the FATP genes being more highly expressed in the MITF^High^ Transient or Melanocytic phenotype cells, and *SLC27A2* and *SLC27A6* being expressed predominantly in a few MITF^Low^ undifferentiated or Neural Crest-like cell lines. *CD36* was expressed to high levels in only 3 lines. These results, together with the observations that lipofermata predominantly affected oleic uptake in MITF^High^ lines, raised the possibility that long chain fatty acid uptake in MITF^Low^ melanoma cells is largely FATP-independent.

One candidate mechanism for fatty acid uptake is via caveolae, cholesterol and sphingolipid-rich lipid rafts containing the integral membrane protein caveolin. Caveolae have been reported to be important for fatty acid transport in a number of cell types including adipocytes (Mattern et al., 2009; Pohl et al., 2004; Pohl et al., 2005; Ring et al., 2006). Remarkably, using the same panel of 53 melanoma cell lines in which different phenotypes can be distinguished using *MITF*, *SOX10*, and *SOX9* as markers (Figure 2E), revealed that significant expression of *CAV1* mRNA encoding *Caveolin 1,* a key component of caveolae in plasma membranes (Conde-Perez et al., 2015), was largely restricted to the MITF^Low^ undifferentiated and neural crest-like melanoma cell lines that also express *AXL*, a hallmark of melanoma invasion and therapy resistance (Konieczkowski et al., 2014; Muller *et al*., 2014). Similar results were obtained using our in-house panel of lines where RNA-seq (Figure 2F) and western blotting (Figure 2G) also confirmed that CAV1 expression was largely restricted to the 4 MITF^Low^/AXL^High^ cell lines. Notably, the correlation between AXL and CAV1 expression was not restricted to melanoma cells but was recapitulated across cancer types in the Cancer Cell Line Encyclopedia (Figure 2H).

To assess the impact of inhibiting caveolae-dependent endocytosis on fatty acid uptake by melanoma cells, we treated cells with oleic acid and used Nystatin, a sterol-binding compound that disassembles caveolae in the membrane. As anticipated the uptake of oleic acid by the MITF^High^ (501mel and IGR37) cell lines was unaffected by Nystatin (Figure 2I, J). By contrast, the accumulation of lipid droplets was reduced by Nystatin in both the MITF^Low^ IGR39 and WM793 cells. Collectively the results indicate that while FATPs are used by the lipofermata-sensitive MITF^High^ melanoma cells, MITF^Low^/AXL^High^ cells may use alternative mechanisms to take up oleic acid, including via caveolae. The observation that undifferentiated melanoma cells use a FATP-independent mechanism for fatty acid uptake is important since blocking FATP activity has been proposed as a potential anti-cancer therapy.

### Oleic acid triggers phenotype-specific translocation of a β-catenin-caveolin complex

To examine the molecular consequences of oleic acid uptake, we focused on the undifferentiated MITF^Low^ melanoma cells that can induce lipolysis in human adipose tissue explants and import oleic acid in a lipofermata-independent fashion. Since CAV1 expression is largely restricted to MITF^Low^/AXL^High^ melanoma cells we performed immunofluorescence to detect CAV1 in the IGR39 cell line used as a model for the BRAFi-resistant undifferentiated phenotype. Under control conditions, CAV1 was excluded from nuclei, using Lamin B as a marker for the nuclear periphery, and was found primarily in the cytoplasm or associated with the plasma membrane as expected (Figure 3A). By contrast, exposure to oleic acid (OA) led to increased CAV1 expression and a proportion of the protein exhibiting a nuclear localization. This observation was confirmed by Western blotting of fractionated cell extracts (Figure 3B) that showed OA could induce elevated levels of cytoplasmic CAV1 as well as an increase in nuclear CAV1. Nuclear accumulation of a proportion of CAV1, and its interaction with the inner nuclear membrane protein emerin, has been noted previously(Chretien et al., 2008; Sanna et al., 2007), though the trigger for nuclear localization of CAV1 and its consequences had not previously been deciphered.

**Figure 3.**
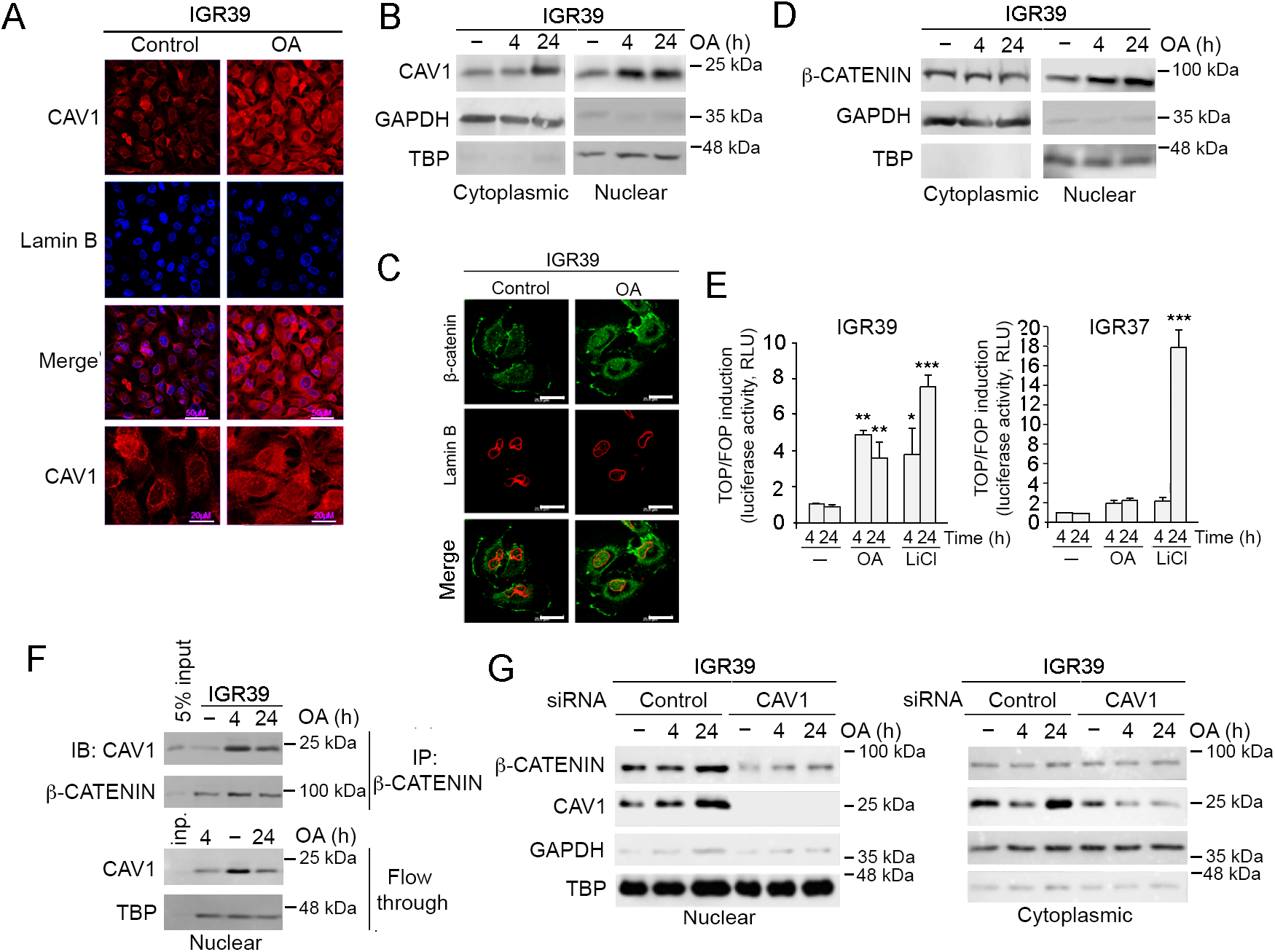
Oleic acid induces β-catenin-CAV1 interaction and nuclear localization. **A**. Immunofluorescence of IGR39 cells using antibodies against indicated proteins in cells exposed to 100 μM oleic (OA). Scale bars indicate 50 µm for top panels and 20 µm for bottom 3 panels. **B**. Western blots for indicated proteins from fractionated IGR39 cells treated or not with 100 μM OA for over time. GAPDH and TBP were used as markers of the cytoplasmic and nuclear fractions**. C**. Immunofluorescence of IGR39 cells for indicated proteins after exposure or not to 100 μM OA for 6 h. Scale bars indicate 25 µm. **D.** Western blot of fractionated IGR39 cells treated with oleic acid for indicated times. **E.** Luciferase reporter assays in indicated cell lines showing the ratio of luciferase activity from the TOP:FOP-flash β-catenin activity reporters. n= 3, error bars indicate SEM ** *P* <0.01, *** *P* <0.001. One-Way ANOVA Statistical test. **F**. Western blot for indicated proteins immunoprecipitated from IGR39 nuclear extract using anti-β-catenin antibody from cells treated or not with OA 100 μM for indicated times. Blot shows both immunoprecipitated (IP) and flow through (FT). **G**. Western blots for indicated proteins from fractionated IGR39 cells (nuclear upper panels; cytoplasmic, lower panels) treated or not with 100 μM OA for indicated times and transfected with either control or CAV1-specific siRNA.

CAV1 can interact with β-catenin (Conde-Perez *et al*., 2015), whose activity is primarily promoted by WNT signaling (Clevers, 2006), together with glucose-driven β-catenin acetylation that promotes its nuclear accumulation and regulation of its target genes (Chocarro-Calvo et al., 2013). β-catenin plays a key role in the melanocyte lineage by driving the genesis of melanoblasts in the neural crest (Sommer, 2011), activating melanocyte stem cells (Rabbani et al., 2011), suppressing melanoma senescence (Delmas et al., 2007), and promoting melanoma proliferation (Widlund et al., 2002) and metastasis (Damsky et al., 2011). The cytoplasmic interaction between β-catenin and CAV1 is inhibited by PTEN, a suppressor of PI3K, (Conde-Perez *et al*., 2015), that is mutated in IGR39 cells. Therefore, we asked whether oleic acid could control CAV1-β-catenin interaction to mediate their nuclear translocation.

Significantly, 4 h after addition of oleic acid we noted an increase in β-catenin nuclear localization in IGR39 cells both by immunofluorescence (Figure 3C), and by Western blotting of fractionated extracts using GAPDH and TBP as markers of the cytoplasmic and nuclear fractions respectively (Figure 3D). The ability of OA to promote nuclear localization of β-catenin in the MITF^Low^ IGR39 cell line was similar to that of LiCl (Supplemental Figure S2A), that prevents β-catenin degradation by inhibiting GSK3. Significantly, in CAV1-negative IGR37 cells, OA had no effect on β-catenin nuclear accumulation, whereas LiCl both increased β-catenin levels and nuclear location (Supplemental Figure S2B). The results obtained using immunofluorescence of IGR37 cells were confirmed by western blotting of fractionated cell extracts (Supplemental Figure S2C). Importantly, in IGR39 cells LiCl induced the inhibitory GSK3β phosphorylation on S9, but oleic acid did not (Supplemental Figure S2D), indicating that the mechanism underlying nuclear localization of β-catenin in response to oleic acid was not mediated by inhibition of GSK3β. Nor did we detect any change in phosphorylation of Y279 or Y216 on GSK3α and GSK3β respectively, modifications that facilitate activation of the WNT-β-catenin pathway (Gao et al., 2015). We also noted no significant change in β-catenin mRNA levels in response to oleic acid (Supplemental Figure S2E).

Consistent with OA triggering nuclear localization of β-catenin, exposure of IGR39 cells to oleic acid for 4 h led to a 5-fold transcriptional activation of the SuperTOP-flash reporter (Veeman et al., 2003) (Figure 3E, left panel) containing canonical β-catenin-LEF/TCF response elements, compared to the non-responsive mutant variant (SuperFOP-flash). As a positive control we used LiCl, that at 24 h increased the β-catenin-responsive TOP -flash reporter by almost 4-fold compared to the non-responsive FOP-flash reporter. Unlike the IGR39 cells, using the MITF^High^ IGR37 cell line we failed to see any significant effect of oleic acid on the TOP-flash reporter at 4 h or 24 h, while a robust activation of the reporter was noted 24 h after addition of LiCl (Figure 3E right panel). Thus, the transcriptional response of cells to oleic acid was phenotype specific.

We next asked whether oleic acid could affect β-catenin nuclear accumulation via promoting an association with CAV1. Co-immunoprecipitation of nuclear extracts using an anti-β-catenin antibody revealed that oleic acid led to increased interaction between β-catenin and CAV1 in the nuclear fraction (Figure 3F). Significantly, CAV1-specific siRNA prevented oleic acid-mediated nuclear accumulation of β-catenin (Figure 3G). Collectively these data suggest that oleic acid triggers formation of a β-catenin-CAV1 complex, and that CAV1 is necessary for oleic acid-induced nuclear translocation and activity of β-catenin. Since CAV1 is only expressed in the MITF^Low^/AXL^High^ undifferentiated cells, these observations uncover a phenotype specific molecular response to oleic acid.

### Oleic acid-mediated activation of SRC promotes CAV1 and β-catenin nuclear localization

Transcriptional activity of β-catenin requires its phosphorylation by SRC at Y333 (Yang et al., 2011). Whether oleic acid could induce β-catenin phosphorylation through SRC activation in IGR39 cells is unknown. To investigate this, cells were treated with oleic acid and β-catenin phosphorylation probed by western blotting using a pY333 antibody. The results revealed that oleic acid induced robust phosphorylation of β-catenin at Y333 (Figure 4A). We also noted that oleic acid induced phosphorylation of CAV1 at Y14 (Figure 4B), also a known SRC phosphorylation site (Li et al., 1996) previously implicated in melanoma cell invasion (Ortiz et al., 2016), as well as increasing CAV1 levels. These data suggest that oleic acid may induce activation of SRC in IGR39 melanoma cells. We therefore assessed the effect of oleic acid over time on both phosphorylation of CAV1 Y14, and on SRC phosphorylation at Y416 that stimulates SRC kinase activity by stabilizing its activation loop. pY416 SRC is frequently used as a surrogate measure of SRC activity (Kmiecik et al., 1988). The results, (Figure 4C) revealed that phosphorylation of CAV1 Y14 increases over time following exposure of cells to oleic acid, starting at 30 min to 1 h. Similarly, within 30 minutes oleic acid induces the activating Y416 phosphorylation of SRC that is maintained over 24 h. We further confirmed that oleic acid mediates activation of SRC by examining phosphorylation of STAT3 Y705, another well-characterized SRC target (Cao et al., 1996). The results (Figure 4C, lower panels) revealed that the increase in SRC Y416 over time induced by oleic acid was paralleled by elevated STAT3 Y705 phosphorylation. Significantly, the activation of SRC by oleic acid was reproduced in an alternative MITF^Low^ undifferentiated melanoma cell line, WM793 (Figure 4D). By contrast, no SRC activation was detected in the MITF^High^ line IGR37. Consistent with SRC phosphorylating CAV1, treatment of IGR39 cells with the SRC family inhibitor PP2 abolished the phosphorylation of CAV1 at Y14 (Figure 4E) as well as reducing CAV1 levels.

**Figure 4.**
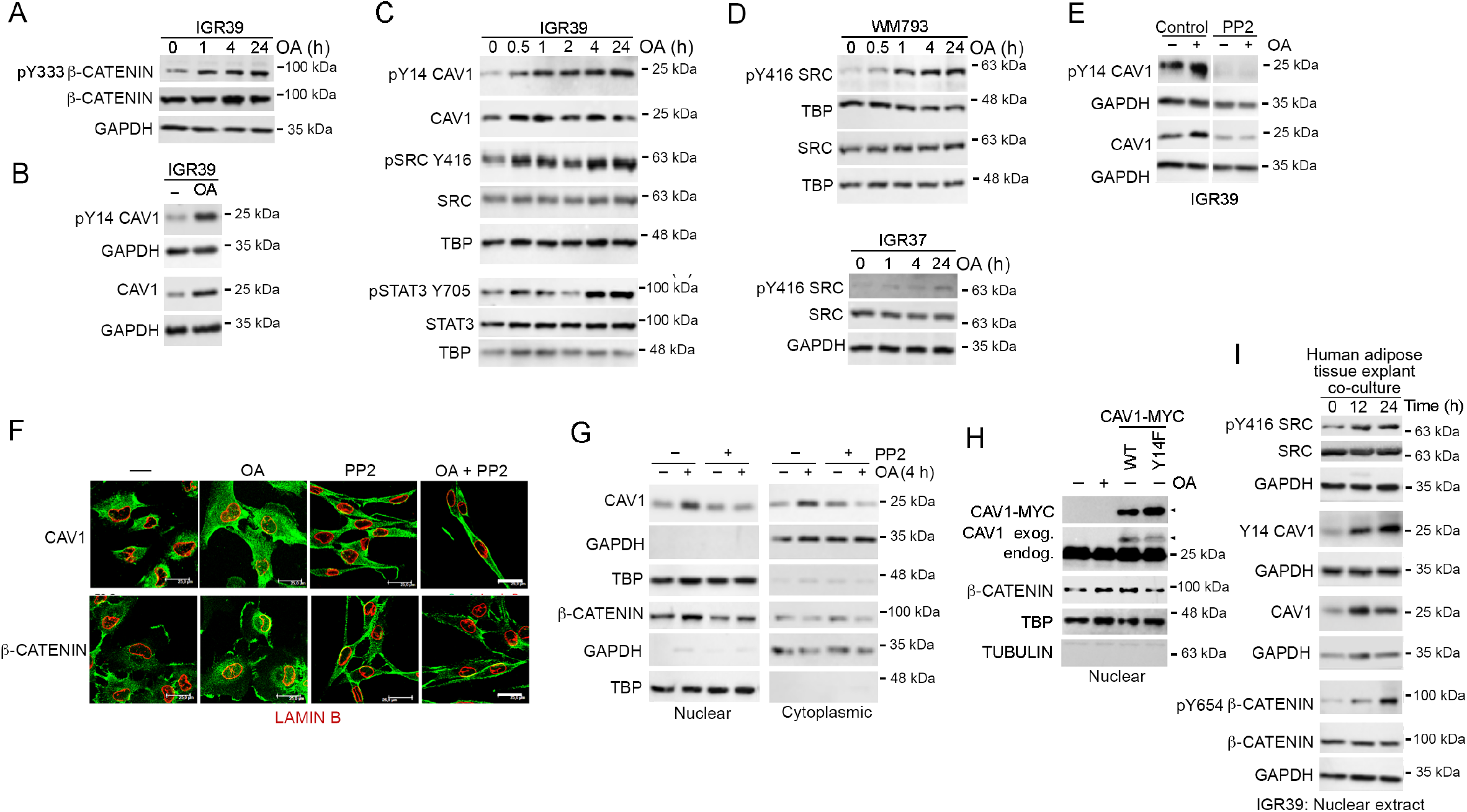
OA activates SRC in MITF^Low^ cells. **A-E**. Western blots for indicated proteins treated with 100 μM (OA). **E.** IGR39 cells were treated 12 h with 10 µM PP2 +/- 10 µM (OA) for another 4 h before being analysed by western blot. **F.** Immunofluorescence for indicated proteins in IGR39 cells pre-treated for 12 h with 10 µM PP2 and then +/- 10 µM (OA) for a further 4 h. **G**. Western blots of fractionated extract treated 12h with 10 µM PP2 +/- 10 µM (OA) for another 4 h. **H**. Western blots for indicated proteins of nuclear or chromatin fractions from IGR39 cells treated with 100 µM oleic acid (OA) and transfected where indicated with expression vectors for WT or Y14F mutant CAV1. Arrowheads indicate ectopic CAV1 WT or Y14F mutant. **I**. Western blot for indicated proteins of nuclear extracts of IGR39 cells in co-culture with adipose tissue explants.

The requirement of OA-induced SRC activation for CAV1-β-catenin complex formation and nuclear localization was tested by inhibition or depletion of SRC. Immunofluorescence revealed that inhibition of SRC family kinases using PP2 not only blocked the oleic acid-dependent increase in nuclear CAV1, but also prevented the nuclear accumulation of β-catenin (Figure 4F). This result was confirmed using Western blotting (Figure 4G), that showed that the oleic acid-induced nuclear accumulation of both CAV1 and β-catenin was blocked using PP2. Notably overexpression of WT CAV1, but not a Y14F non-phosphorylatable mutant, was able to promote nuclear accumulation of β-catenin, mimicking the effect of oleic acid (Figure 4H).

As these results were obtained using free oleic acid, we next ascertained whether the bi-directional interaction between melanoma cells and human adipose tissue explants could recapitulate the observations. The results revealed that like oleic acid, exposure of IGR39 cells to human adipose tissue explants led to activation and phosphorylation of SRC and downstream phosphorylation of CAV1 Y14 and β-catenin Y654 (Figure 4I).

### Oleic acid activates AXL

Taken together the data so far reveal that undifferentiated MITF^Low^ melanoma cells, but not MITF^High^ cells, induce lipolysis in human adipose tissue explants, and that FATP-independent fatty acid uptake by the melanoma cells triggers a cascade of events including activation of SRC leading to formation and nuclear translocation of a CAV1-β-catenin complex. However, even though both MITF^High^ IGR37 and MITF^Low^ IGR39 cells can take up free oleic acid, SRC is only activated in MITF^Low^ IGR39 cells.

Since SRC can be activated by receptor tyrosine kinase (RTK)-mediated phosphorylation, we considered the possibility that oleic acid-induced SRC phosphorylation would be mediated through activation of an RTK whose expression is restricted to MITF^Low^ cells. One candidate was AXL, a key RTK whose expression has been associated with resistance to therapy in melanoma (Konieczkowski *et al*., 2014; Muller *et al*., 2014) as well as breast and other cancers (Auyez et al., 2021). Importantly, AXL is specifically expressed in MITF^Low^ melanoma cells (Figure 2E-G) and is normally activated by its ligand GAS6 that promotes its dimerization. Instead, we considered the possibility that activation might also be achieved via exposure of cells to oleic acid. We initially asked whether oleic acid would trigger autophosphorylation in trans of Y779, a marker of AXL activation. Western blotting of the MITF^Low^, AXL^High^ WM793 melanoma cell line revealed that oleic acid induced elevated AXL Y779 phosphorylation, indicative of AXL activation, starting within 30 min of oleic acid addition (Figure 5A). In both IGR39 and in WM793 cells, activation of AXL was reflected by elevated SRC phosphorylation (Figure 5B). Importantly, the increase in SRC phosphorylation triggered by exposure to oleic acid in both IGR39 or WM793 was blocked by treatment with the AXL inhibitor ONO-7475 (Figure 5C). Inhibition of AXL using ONO-7475, or a second inhibitor R428 (Bemcetinib), also prevented the induction of phosphorylation of CAV1 Y14 by Oleic acid (Figure 5D). Thus oleic acid triggers a cascade of downstream events leading to nuclear localization of a CAV1-β-catenin complex (Figure 5E).

**Figure 5.**
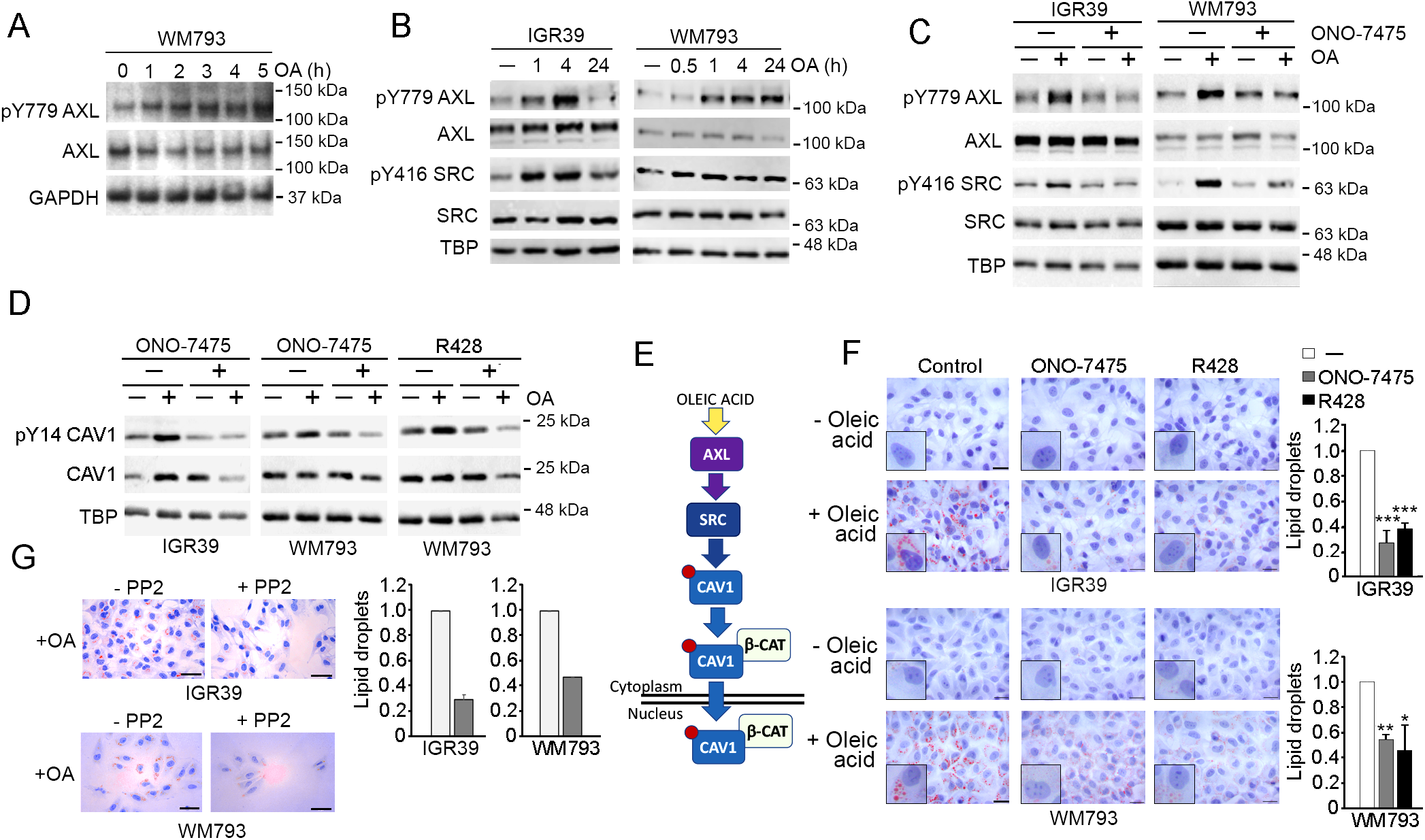
Oleic acid activates AXL. **A-D.** Western blots of indicated cell lines treated with 100 μM oleic acid for 4 h unless indicated, and/or pre-treated with AXL inhibitors ONO-7475 (0.5 µM for IGR39 or 0.1 µM for WM793) or R428 (0.5 µM) for 2 h prior to exposure to OA. **E**. Schematic summarising signalling downstream from oleic acid. **F**. Oil Red O staining of IGR39 and WM793 cells exposed to ONO-7475 (0.5 µM) or R428 (0.5 µM) for 2h +/- 100 µM OA for 12 h more. AXL inhibitors ONO-7475 (0.5 µM ) or R428 (0.5 µM ). Insets show enlarged examples of typical cells. Scale bars indicate 50 µm. Quantification (right) of lipid droplet accumulation N=3, Error bars indicate SEM * *P* <0.05, ** *P* <0.01***, *P* <0.001. One-Way ANOVA Statistical test.

Intriguingly, treatment of cells with the AXL inhibitors ONO-7475 or R428 decreased oleic acid uptake into both IGR39 and WM793 melanoma cells (Figure 5F), as did inhibition of SRC with PP2 (Figure 5G). This is consistent with oleic acid uptake in MITF^Low^ melanomas being mediated by caveolae (Figure 2I, J) since CAV1 Y14 phosphorylation by SRC is necessary for caveolae-mediated endocytosis (Hau et al., 2019). The results also reveal a potential positive feedback loop between oleic acid uptake and AXL activation.

### Oleic acid-mediated activation of AXL drives melanoma invasiveness

Since AXL and SRC activation may drive invasion, we assessed whether OA could enhance invasion in IGR39 cells. The results obtained using a Matrigel transwell assay revealed that OA induced a robust increase in invasion in MITF^Low^ IGR39 cells, but not in MITbodipyF^High^ IGR37 cells (Figure 6A). Similar results were obtained using human adipose tissue explants that in co-culture induced invasion in IGR39, but not IGR37 cells (Figure 6B). siRNA-mediated depletion of either CAV1 or β-catenin (Figure 6C) inhibited OA-induced invasion (Figure 6D). Importantly, the OA-mediated increase in invasion by IGR39 cells was also prevented using two different AXL inhibitors, ONO-75475 or R418 (Figure 6E).

**Figure 6.**
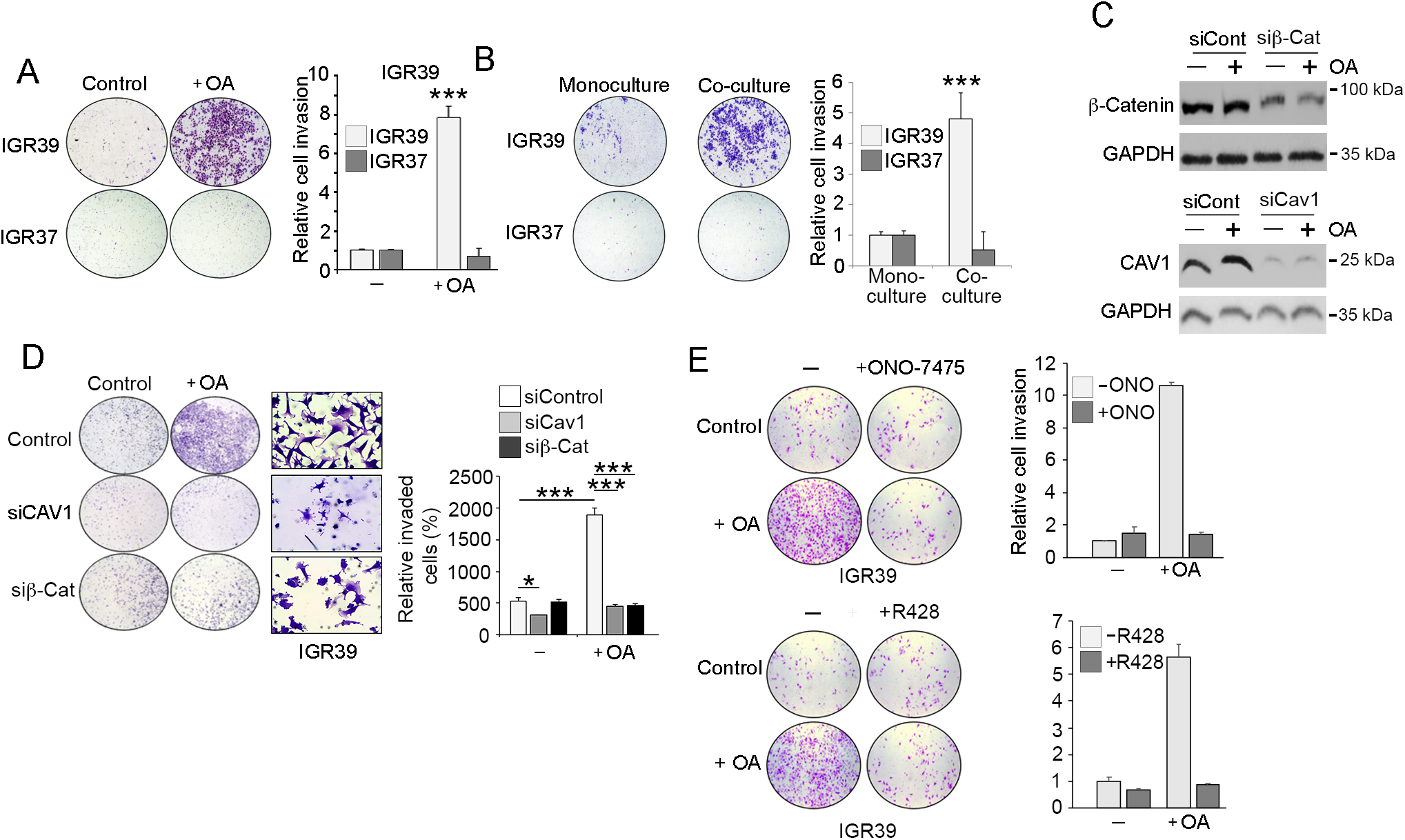
Oleic acid induces AXL-dependent invasion in melanoma. **A,B,D,E**, Matrigel transwell invasion assays showing invading cells stained with crystal violet. Where indicated cells were treated with 100 μM oleic acid, and/or transfected with control or siRNAs targeting CAV1 or β-catenin, or co-cultured with adipose tissue explants, or treated with AXL inhibitors ONO-7475 (0.5 µM) or R428 (0.5 µM). Assays were performed for 22 h. Quantification from n≥ 3 biological replicates; error bars = SEM; * =P< 0.5, ** =P< 0.1, *** = p< 0.01. One-Way ANOVA Statistical test. **C.** Western blot showing depletion of CAV1 or β-catenin after transfection with indicated control or specific siRNAs. GAPDH was used as a loading control.

## Discussion

It is now widely appreciated that tumors contain multiple phenotypically distinct populations of cancer cells that differ in their biological properties, some of which contribute to therapy resistance and metastatic dissemination. Yet whether specific phenotypic states might mediate unique bi-directional interactions with the microenvironment is not well understood. Here we reveal a previously unsuspected ability of oleic acid, a widely available microenvironmental fatty acid, to activate AXL, a phenotype-restricted RTK implicated in therapy resistance. OA-driven activation of AXL leads to downstream SRC signalling, nuclear localization of a complex between CAV1 and β-catenin, a key melanocyte and melanoma effector protein, and increased invasion.

In many cancer types AXL expression has been associated with an epithelial to mesenchymal (EMT) transition and invasiveness (Asiedu et al., 2014; Boshuizen et al., 2018; Shao et al., 2023; Wang et al., 2016) and is linked to worse prognosis and resistance to therapies in a wide range of cancers including prostate (Bansal et al., 2015), breast (Creedon et al., 2014), lung (Zhang et al., 2012), renal (Zhou et al., 2016), head and neck (Elkabets et al., 2015) and neuroblastoma (Debruyne et al., 2016). Consequently, AXL has attracted increasing attention as a candidate for therapy (Auyez *et al*., 2021; Colavito, 2020; Gay et al., 2017). In melanoma, high AXL is specifically expressed in undifferentiated and invasive phenotype phenotype cells and is commonly associated with therapy-resistance (Konieczkowski *et al*., 2014; Muller *et al*., 2014; Rambow *et al*., 2018) as well as resistance to immune checkpoint inhibition (Hugo et al., 2016). As a consequence, targeting AXL can eliminate specific subpopulations of melanoma cells and can cooperate with inhibition of the MAPK pathway (Boshuizen *et al*., 2018). Importantly, rather than AXL expression simply acting as a marker of therapy resistance and EMT, in pre-clinical models targeting AXL can sensitize tumors to therapies (Auyez *et al*., 2021). This suggests that to promote therapy resistance and to exert its pro-survival and pro-metastasis effects, AXL should be activated by its ligand GAS6, which is widely expressed by cancer cells, stromal cells and infiltrating immune cells (Auyez *et al*., 2021). Alternatively, as we show here, AXL can also be activated by oleic acid, triggering downstream activation of SRC. Our results therefore highlight a way in which the availability of a key nutrient can differentially impact distinct classes of melanoma cells via ligand-independent activation of an RTK with major consequences for disease progression. This is important as AXL is expressed on invasive and undifferentiated phenotype cells that may escape tumors to enter the lymphatic or blood vessels, or invade other tissues where GAS6 may not be abundant. Under these circumstances, an alternative pathway to activating AXL will provide a critical selective advantage. For example, oleic acid that is the major circulating free fatty acid in tumor associated lymph (Morfoisse et al., 2021), including in melanoma (Ubellacker *et al*., 2020), and which can be released from adipocytes in proximity to melanoma cells (Zhang *et al*., 2018), may promote survival and maintenance of an invasive state via AXL signaling. Moreover, as activation of SRC is a pro-survival signal triggered by integrin-interactions with extracellular matrix that has been linked to BRAFi-resistance in vivo (Hirata et al., 2015), it is plausible that the ability of oleic acid to promote SRC activation via AXL provide a degree of adhesion mimicry that would suppress cell death in non-attached, metastasizing cancer cells. In this respect, the expression of AXL on therapy resistant cells in a wide variety of cancers known to invade or metastasize to adipose tissue (Hoy *et al*., 2021) or via the lymphatic system may provide a key pro-survival mechanism that extends well beyond the ability of some fatty acids, such as palmitate (Altea-Manzano et al., 2023), to promote proliferation via fatty acid oxidation.

In addition to fatty acid uptake from lymph or through melanoma induced lipolysis in adjacent adipocytes, the activation of AXL by oleic acid may also have implications for our understanding of the impact on cancer progression of obesity, a known risk factor for many cancers including melanoma (García-Jiménez et al., 2016). Obesity is associated with elevated plasma free fatty acid (FFA) levels (Henderson, 2021). As a consequence, the elevated availability of FFAs in obese individuals will increase the probability of activation of AXL and its downstream signaling in MITF^Low^ melanoma cells, promoting survival, increased invasiveness and metastatic dissemination as well as therapy resistance. Consistent with this, mice fed a high fat diet exhibit increased melanoma progression associated with elevated Cav1 as well as pCav1 (Pandey et al., 2012).

Beyond activation of AXL our results provide three additional and unanticipated insights. First, we show that unexpectedly, only MITF^Low^, undifferentiated melanoma cells, and not MITF^High^ cells, are competent to induce lipolysis in human adipose tissue explants; second, in contrast to MITF^High^ cells, uptake of oleic acid by MITF^Low^ melanoma cells does not appear to involve lipofermata-sensitive FATPs or CD36; and third, our results highlight the phenotype specific effect of oleic acid in promoting, via activation of AXL and SRC, nuclear translocation of a CAV1-β-catenin complex. These observations have profound implications for our understanding of how bidirectional interactions with adipocytes in vivo may shape disease progression.

The observation that MITF^Low^, but not MITF^High^, melanoma cells can induce lipolysis in, and lipid uptake from, human adipose explants was unexpected. However, this observation is entirely in keeping with the hypothesis that MITF^Low^ cells are invasive because they lack, or fail to sense, key nutrients that are needed to support log-term survival (García-Jiménez and Goding, 2019). For example, as MITF activates transcription of the key fatty acid desaturase SCD, MITF^Low^ cells express low levels of SCD and exhibit an increased saturated to mono-unsaturated fatty acid ratio, an imbalance that can stabilise an invasive state (Vivas-Garcia *et al*., 2020). Therefore, it makes sense for invasive MITF^Low^ cells to try to redress this imbalance by acquiring fatty acids from adipocytes. However, while lipid release may be triggered by invasive MITF^Low^ cells, it does not mean they will revert to a proliferative state which would require both the carbon provided by fatty acids, but also nitrogen (amino acids) to fuel protein synthesis. Thus, lipolysis and fatty acid uptake may increase invasion to enable cells to maintain their search for a niche suitable to promote proliferation.

Why MITF^High^ phenotype cells use FATPs to take up fatty acids, while uptake in MITF^Low^ cells is FATP independent is unclear. As FATPs have been found in species from bacteria and yeast to man (Doege and Stahl, 2006), it might have been expected that all cells would use FATPs to transport fatty acids. One possible explanation for this phenotype-specific difference, is that undifferentiated MITF^Low^ cells tend to be slow cycling and would have reduced demands for energy and the building blocks, especially membranes, required to generate daughter cells compared to more rapidly dividing MITF^High^ cells. It is plausible therefore that the need for long-chain fatty acids (LCFAs) by slow cycling MITF^Low^ cells can be met by non-FATP-mediated transport, but in proliferative MITF^High^ cells the high demand for LCFAs requires efficient transport by FATPs. This hypothesis would be consistent with the recent observation that the melanocytic state, associated with MITF^High^-driven differentiation, is characterized by increased uptake of oleic acid, and accumulation of lipid droplets (Lumaquin-Yin *et al*., 2023). Importantly, the observation that MITF^Low^ cells use a FATP-independent mechanism for fatty acid uptake has significant implications for the therapeutic use of anti-cancer strategies that propose the use of FATP or CD36 inhibitors as an effective anti-cancer strategy (Alicea *et al*., 2020; Pascual *et al*., 2016; Zhang *et al*., 2018); MITF^Low^ cells would be resistant to such inhibitors as they are to many other drugs including BRAF inhibitors.

Notably, previous work has failed to identify a nuclear localization signal in β-catenin, meaning that its nuclear localization presumably arises through regulated interaction with cofactors. Here, we reveal that oleic acid activates a novel AXL, SRC and CAV1-dependent mechanism that can promote nuclear localization of β-catenin independent of addition of WNT, the canonical trigger for β-catenin nuclear accumulation. Although previous work has shown cytoplasmic β-catenin-CAV1 interaction in the absence of PTEN (Conde-Perez *et al*., 2015), neither the ability of this complex to translocate to the nucleus, nor the regulation of β-catenin-CAV1 complex formation by an oleic acid/AXL/SRC axis, was known. Significantly, the ability of OA to drive nuclear translocation of β-catenin is restricted to MITF^Low^ melanoma cells, since only the neural crest-like and undifferentiated phenotypes express CAV1 and AXL whose expression we find is highly correlated in melanoma cells as well as in other cancer types.

In summary, we reveal here a phenotype specific ability of de-differentiated cancer cells to induce fatty acid release from human adipocytes that are then taken up via a non-FATP mechanism to promote activation of AXL, SRC and β-catenin signalling to drive increased invasion and metastatic potential.

## KEY RESOURCES TABLE

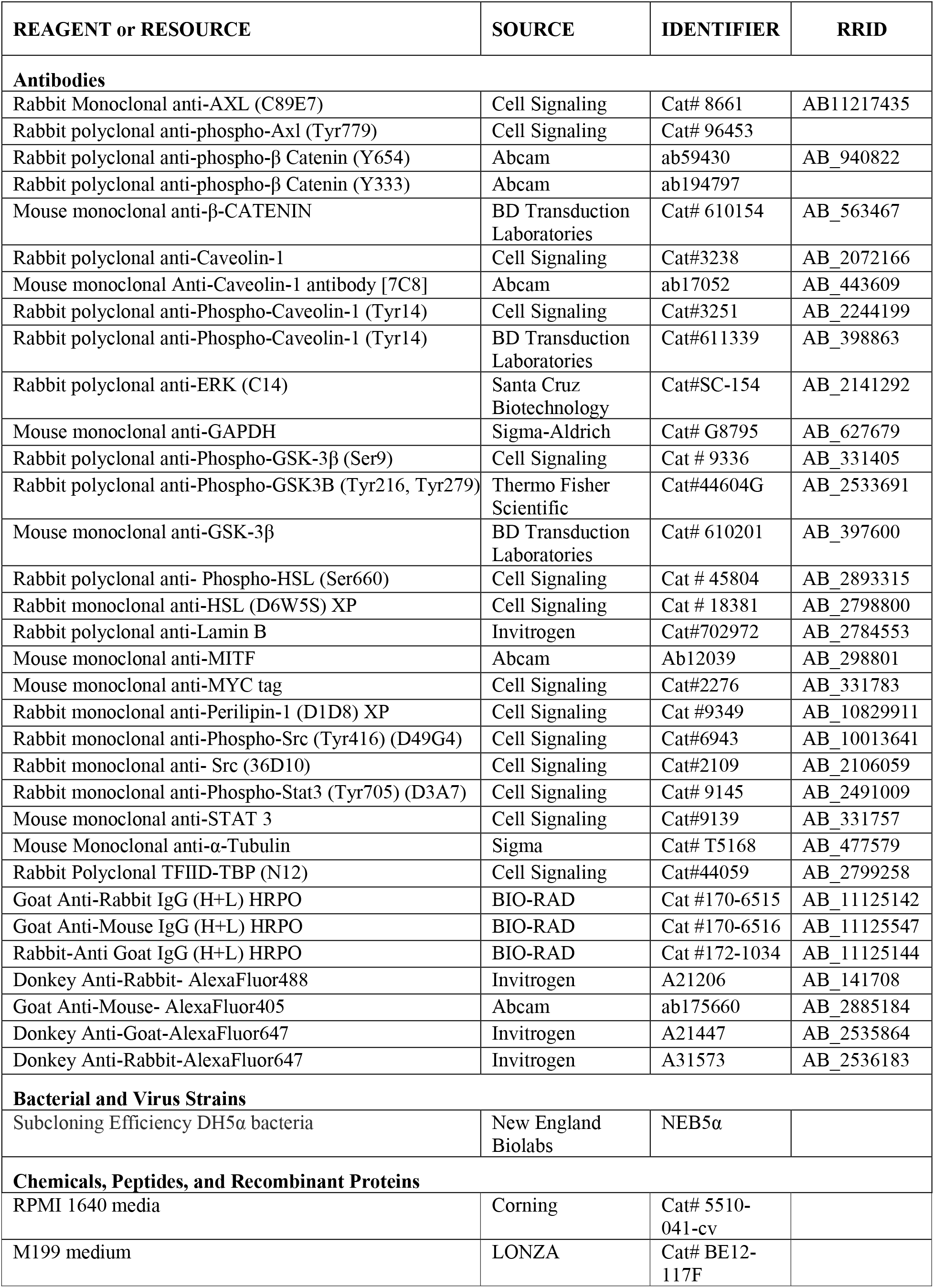
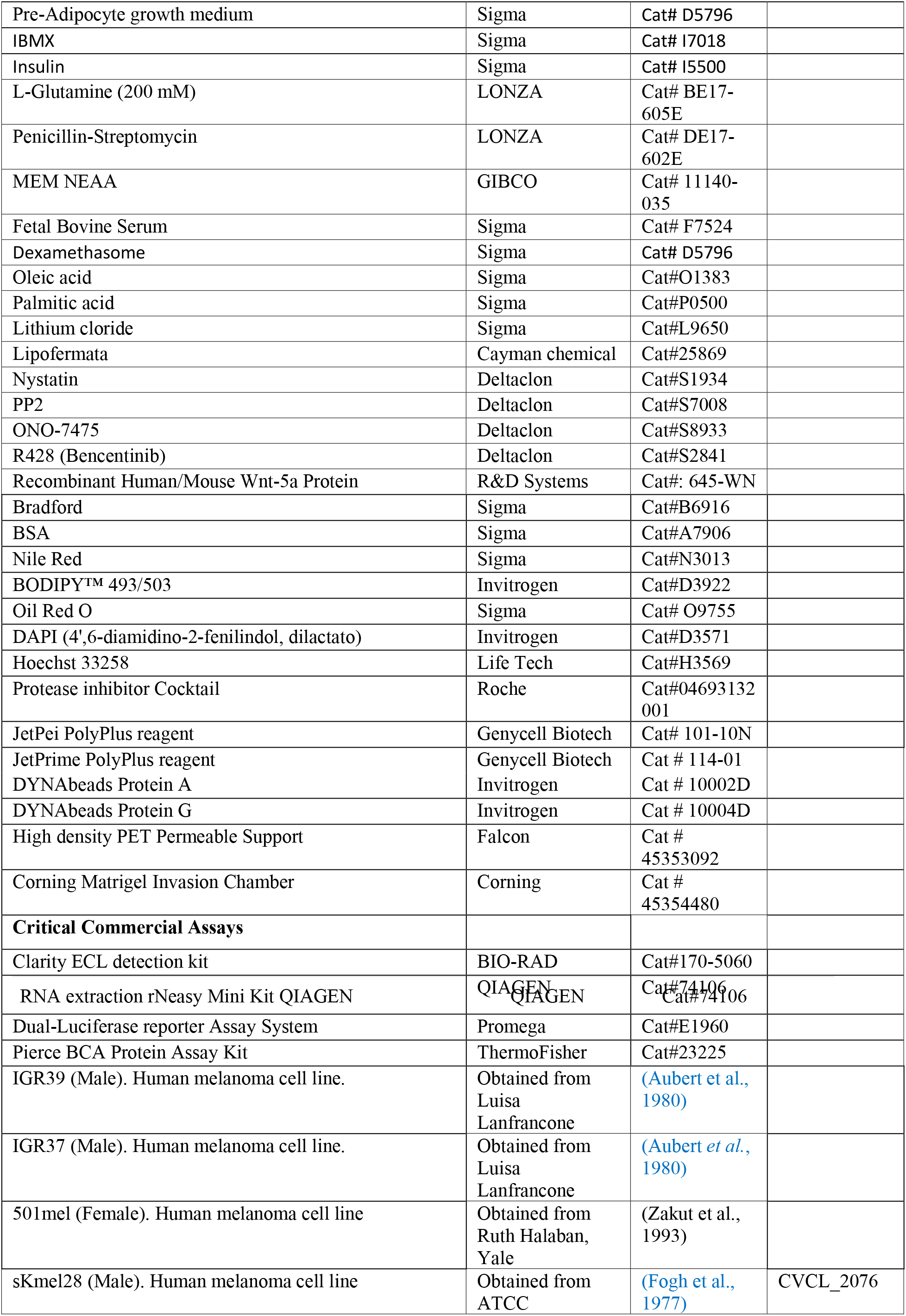
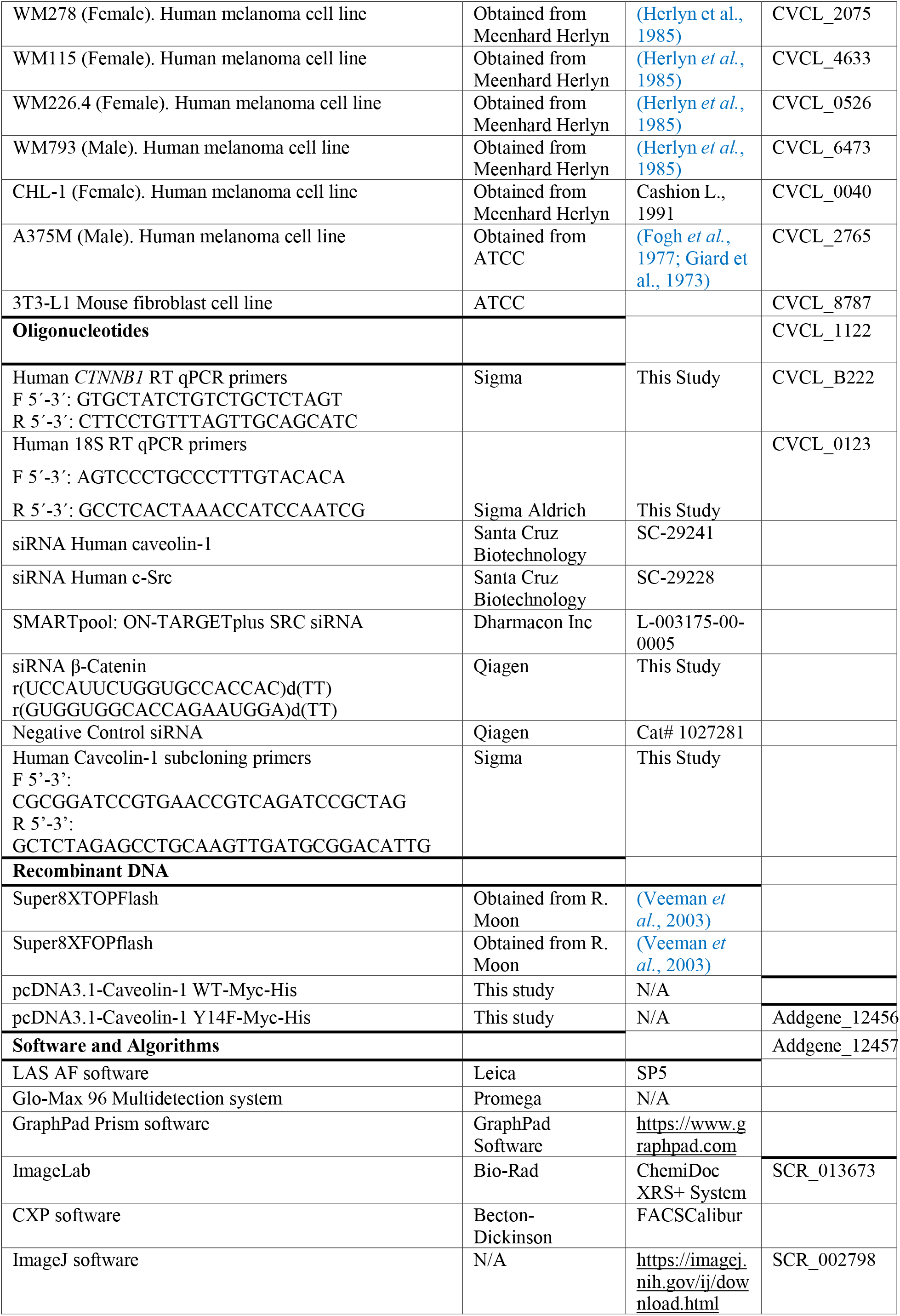
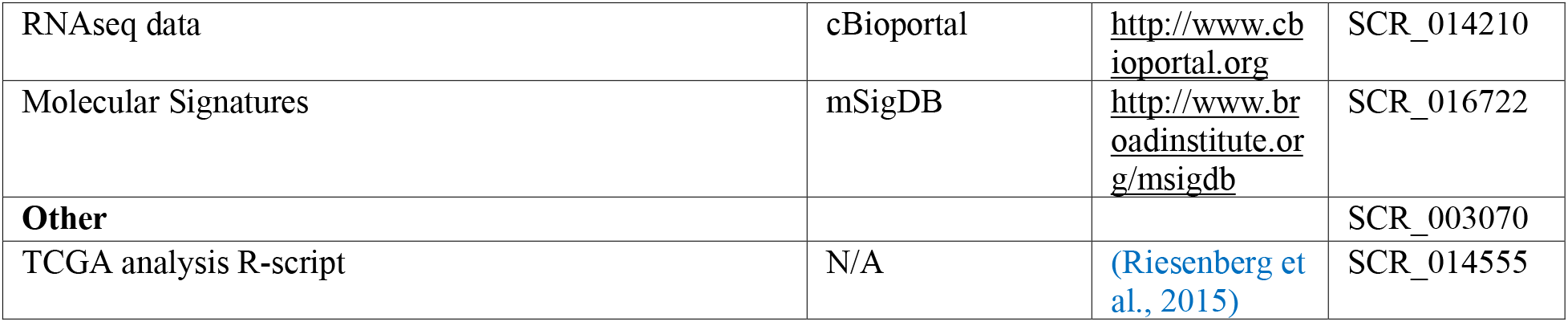

## METHODS

### Cell lines

All human melanoma cell lines were cultured in RPMI supplemented with 10% fetal bovine serum (FBS) plus 1% penicillin–streptomycin and maintained in 5% CO^2^ environment at 37°C. All the cell lines were tested monthly for mycoplasma and subject to authentication by short tandem repeat (STR) profiling.

### RNAi gene silencing

Cells plated in six well plates at 50% confluence were transfected with siRNA using JetPRIME Polyplus reagent (Genycell Biotech, Santa Fe, Granada, Spain), following the manufacture’’s instructions. After 2 days cells were treated and collected to perform invasion assay or to be analysed by western blot. Specific siRNA oligonucleotides for β-catenin and control siRNA were obtained from Qiagen, Caveolin-1, and SRC siRNA from Santa Cruz Biotechnology and SRC siRNA Dharmacon Inc.

### Plasmid subcloning

Caveolin-1 WT or Y14F were subcloned from CAV1-mRFP plasmid (Tagawa et al., 2005) by PCR amplification with the indicated primers (Key Resources table) followed by a digestion with Bam HI and Xba I enzymes (New England Biolabs) for 37°C 2 h. The fragments were gel-purified with a kit (Qiagen, Crawley, UK) were cloned into pcDNA3.1-myc-His plasmid (Promega). For sequencing we used a sequence analyser (ABI Prism 3100 Avant; Applied Biosystems, Alcobendas, Madrid, Spain. The correct introduction of the fragment was evaluated by plasmid sequencing using the BigDye Cycle Sequencing Kit (Applied Biosystems).

### Transient transfections

Cells were seeded in plates at 50% confluence and transfected using JetPei PolyPlus reagent (Genycell Biotech, Santa Fe, Granada, Spain), following the manufacturer’s instructions. After 36 h cells were treated as indicated and collected to analyse by western blot or immunofluorescence.

### Preparation of cell extracts

#### Whole cell extracts

Cells were washed with iced PBS before extract preparation and scraped in RIPA buffer (10 mM Tris.HCl pH 7.4, 5 mM EDTA, 5 mM EGTA, 1% Triton X100, 10 mM Na^4^P^2^O^7^, pH 7.4, 10 mM NaF, 130 mM NaCl, 0.1% SDS, 0,5% Na-deoxycholate). After 5 min on ice, cells were pelleted (12,000 rpm for 5 min, 4°C) and the supernatant was directly used as whole cell extract or frozen at -80 °C.

#### Fractionated cell extracts

After washing as before, cells were scraped in hypotonic buffer (20 mM Hepes, pH 8.0, 10 mM KCl, 0.15 mM EDTA, 0.15 mM EGTA, 0.05% NP40, and protease inhibitors) and swollen on ice for 10 min before adding 1:2 vol of sucrose buffer (50 mM Hepes, pH 8.0, 0.25 mM EDTA, 10 mM KCl, 70% sucrose). Lysates were fractionated (5,000 rpm for 5 min at 4°C) to obtain the cytoplasmic fraction in the supernatant. Nuclear pellets were further washed twice with washing buffer (20 mM Hepes, pH 8.0, 50 mM NaCl, MgCl^2^ 1.5 mM, 0.25 mM EDTA, 0.15 mM EGTA, 25% glycerol and protease inhibitors), pelleted at 5,000 rpm, 5 min at 4°C and resuspended in nuclear extraction buffer (20 mM Hepes, pH 8.0, 450 mM NaCl, MgCl^2^ 1.5 mM, 0.25 mM EDTA, 0.15 mM EGTA, 0.05% NP40, 25% glycerol and protease inhibitors) before centrifugation at 12,000 rpm for 5 min at 4°C to pellet and discard cell debris. The supernatants were used as nuclear fractions.

### Immunoprecipitation

Whole cell extracts were obtained in the same buffer without 0.1% SDS and 0,5% Na-deoxycholate. For immunoprecipitation from fractionated extracts the hypotonic buffer was modified by adding 100 mM NaCl and 0.1% NP40. For immunocomplex formation, protein A/G-coated magnetic beads were washed 3 times with the extraction buffer before coating with the primary antibody for 2 h at 4 °C in a rotating wheel, followed by 2 washes with the same buffer to eliminates unbound antibody and then extracts are added O/N at 4°C in the rotating wheel. Immunocomplexes were washed twice and used for western blotting.

### Western blot

Protein lysates were subjected to 7.5 or 10% polyacrylamide SDS-PAGE. Proteins were transferred onto polyvinylidene difluoride membranes. Membranes were blocked with 5% nonfat milk or bovine serum albumin in TBS containing 0.1% Tween 20 and probed with the appropriate primary antibodies (see Key Resources Table) overnight at 4°C. The specific bands were analyzed using ChemiDoc Imaging Systems (Bio-Rad).

### Immunofluorescence microscopy

Cells in cover slips were washed three times and fixed with 4% paraformaldehyde in PBS, pH 7.4 for 10 min; washed again; permeabilized (PBS pH7.4, 0.5% Triton X-100, 0.2% BSA) for 5 min; blocked (PBS pH7,4, 0,05% Triton X-100, 5% BSA) for 1 h at room temperature; incubated with primary antibody over night at 4°C, washed three times for 5 min and incubated with the secondary antibody for 1 h at room temperature. For lipid staining after antibody incubation, cells were washed twice with PBS and stained with a solution containing 5 µg/mL BODIPY or Nile Red. After 30 min incubation at room temperature, cells were washed twice with PBS. Slides were mounted and images were acquired using a SP5 confocal microscope (Leica) with a 63x objective.

### Luciferase reporter assay TOP/FOP reporter Luciferase activity

Melanoma cells seeded in 24-weel plates at 50% confluence were co-transfected with 125ng of the indicated promoter reporter and 25ng of Renilla luciferase construct, for normalization of transfection efficiency, using JetPei PolyPlus reagent (Genycell Biotech, Santa Fe, Granada, Spain), following the manufacturer’s instructions. Forty-eight hours after transfection, cells were treated with 100 µM oleic acid (OA) or 20mM lithium chloride for another 4 or 24 hr. Cells were lysed, and luciferase activity was measured in triplicate using the dual luciferase reporter kit (Promega) and a Glo-Max 96 Multidetector system (Promega). A minimum of 3 experiments were performed per cell line.

### Crystal violet staining and cell density quantification

30,000 cells were plated in a 12-well plate and treated with DMSO, and oleic acid (100 μM) simultaneously over 5-7 days as indicated. Thee medium was replaced every 3 days. Plates were collected at time 0, 24, 72, 96 or 120 h, fixed with 4% PFA and stained with 0.1% crystal violet for 15-30 min. Then they were washed and dried. Crystal violet was resuspended methanol), transferred to p96 plates and analysed by a Spectra FLUOR (Tecan) at 570 nm. Viability was measured in duplicate in four independent experiments.

### 3T3-L1 co-culture experiments

3T3-L1 cells (70,000 cells/well - 12 well plate) were grown to 100% confluency in preadipocyte Growth media (High glucose DMEM (Sigma D5796); 10% Newborn Calf Serum (Sigma 12023C); 1% Penicillin/Streptomycin (Sigma P0781); 1% L-Glutamine (Sigma G7513). When cells reached confluence, the medium was changed, and two days later differentiation was initiated. At day 0, differentiation medium with full induction cocktail was added (High Glucose DMEM (Sigma D5796); 10% FBS; 1% Penicillin/Streptomycin; 1% L-Glutamine; Induction cocktail: 0.5 mM IBMX (Sigma I7018); 1 µM Dexamethasome (Sigma D1881); 5 µg/ml Insulin (Sigma I5500). At Day 3, the medium was changed to differentiation media containing only Insulin (5 µg/ml) for 72 h. At day 6, once mature adipocytes were obtained, the medium was changed to simple differentiation media to preserve their adipocyte differentiated state. Differentiated adipocytes were used for all co-culture experiments.

For lipid transfer experiments, differentiated 3T3-L1 cells were preloaded with 1.5 µM BODIPY-FL-C16 in differentiation media (FBS+insulin) for 4 h, and then washed 2x by using PBS+ 0.1% BSA-free fatty acid. After 24 h, the preloaded adipocytes were washed again with PBS+ 0.1% BSA-free fatty acid, and placed in co-culture with melanoma cells that had been seeded 24 h earlier in the trans-well chambers. The adipocytes were located at the bottom, the melanoma cells on top, and maintained for 24 h in DMEM with 0.1% BSA-free fatty acid and without FBS. After 24 h in co-culture, melanoma cells were washed and fixed with 4% PFA. For picture acquisition, the trans-well membranes were cut, stained with DAPI and mounted in coverslips.

### Adipose tissue and melanoma cell co-culture experiments

Adipose tissue human samples were derived from surgical removal at Hospital Universitario Fundación Alcorcón (HUFA). Immediately after surgery adipose tissue was place into Medium 199 (M199) (Lonza) with penicillin (100 U/l), and streptomycin (100 µg/ml), at room temperature. Blood vessels and connective tissue was removed and uniform-sized tissue pieces of ∼2 mm3 were dissected.

For co-culture experiments 48h before performing the co-culture experiments melanoma cells were seeded at 50% confluence in a six well plate and adipose tissue pieces were cultured with melanoma cell medium. To perform fat-explant-melanoma co-culture experiments high density PET permeable supports with 3.0 µm pore were used (Falcon). The insert was placed in the six well plate where melanoma cells had previously been seeded and 4-5 pieces per well of adipose tissue were placed in the top, 2ml medium of the indicated melanoma cell line was added. 48 hours later the fat and cells were collected and analysed changes in invasion capacity or by WB or immunofluorescence. To test if lipolysis was induced by Wnt5a melanoma IGR37 cells co-cultured with adipose were treated with Wnt5a (50ng/ml) for 4 days. Fat was collected and analysed by WB.

### Oil red O staining

Oil red O stock solution was prepared dissolving 0.5 g of oil red O (Sigma-Aldrich) in 100 ml isopropanol for 1 h at 56°C in a water bath. A working solution was prepared by mixing 30 ml of the Oil red O stock solution with 20 ml of distilled water, allowing to stand for 10 mins, and then filtering. Cells in cover slips were washed three times and fixed with 4% paraformaldehyde in PBS, pH 7.4 for 10 min; briefly washed with running tap water, rinsed with 60% isopropanol, and stained with freshly prepared Oil Red O working solution 15 mins. Then rinsed with 60% isopropanol, nuclei were stained with hematoxylin for 1 min, rinsed with distilled water. Slides were mounted and images were acquired using a Zeiss optical microscope with a 40x objective. Oil red O intensity was quantified using Image J software. For each experiment, 3 different fields were evaluated per slide.

### Matrigel invasion assays

Matrigel invasion assays were performed using invasion chambers from BD Biocoat 8 µm membrane inserts with Matrigel coating. 100,000 cells previously transfected and treated as indicated, or co-cultured with adipose tissue explants were cultured in medium without 0.2% FBS overnight and seeded per insert. Medium with 10% FBS was added to the bottom of the inserts. After 22 h of incubation, chambers were fixed in 3.7% paraformaldehyde for 2 min, washed in phosphate-buffered saline (PBS), stained with 1% crystal violet for 20 min and washed in phosphate-buffered saline (PBS). Cells remaining above the insert membrane were removed by gentle scraping with a sterile cotton swab. Slides were mounted and images were acquired using a Zeiss optical microscope with a 5x objective. Cells were manually counted using ImageJ software. At least three biological replicates were performed.

### RT-qPCR

Total RNA was isolated using the rNeasy minikit (Qiagen) or with Trizol (Invitrogen). cDNA was generated from 1 µg of RNA following the manufacturer’s protocol. Reactions were performed in SYBR Green mix (Go-Taq, Promega) and analyzed using 7500 Fast Real-Time PCR System (Applied Biosystems). Primers for human genes were designed using the Primer Blast application from NCBI (see Key resources table). 18S ribosomal RNA and β-actin primers served as a nonregulated control. Relative expression was calculated using the Ct method, expressed as 2−ΔΔCt (Livak and Schmittgen, 2001). The PCR efficiency was ∼100%.

### Statistical analysis

Results are presented as fold induction, mean ± SEM from at least three biological replicates. Comparisons between two independent groups were performed with Mann-Whitney U-test. Tests for significance between two sample groups were performed with Student’s t test and for multiple comparisons, ANOVA with Bonferroni’s post-test.

### RNA-seq

STR authenticated 21 urofinss genomic service), mycoplasma free (Lonza #LT07-318, #LT07-518 (control) melanoma cell lines were seeded in 6 well tissue cultured dishes in RPMI 1640 (Gibco) supplemented with 10% FBS (Biosera) and 100 U/ml Pen Strep (Gibco) and maintained in 10% CO^2^ humidified chamber. Cells were collected at around 80% confluence and snap-freeze on dry ice before batched processed using rNeasy Mini Kit (QIAGEN #74106) as per supplier instructions and eluted in 50 µl nuclease free water (Invitrogen #10977049). Samples with RIN values ≥9.5 (assessed using using Agilent RNA 6000 Nano Kit (Agilent #5067-1511)) were carried forward for library prep using QuantSeq Forward kit (LEXOGEN #015.96) with 500 ng input material and ERCC ExFold RNA Spike-In Mixes (ThermoFisher #4456739). Sequencing was carried out on a HiSeq4000 (Illumina) by the Oxford Genomics Centre, Wellcome Trust Centre for Human Genetics.

### Bioinformatics

Raw fastq reads for the same samples sequenced across 2 lanes were stitched using UNIX, qCed using fastqc (v0.11.9), adaptor- and poly-A-trimmed using Cutadapt. Processed fastq were mapped against hg38 (GRCh38, 2015) + ERCC STAR index using rna-star (v2.5.1b) with quantMode enabled, allowing for soft-clipping and splicing (min 20 bp). Normalisation and differential gene expression analyses was performed as previously described (Louphrasitthiphol et al., 2020; Louphrasitthiphol et al., 2019; Vivas-Garcia *et al*., 2020) using edgeR glmQLFTest. Heatmaps were visualised using R package pheatmap (v1.0.12).

Gene expression from 53 melanoma cell lines (Tsoi *et al*., 2018) were obtained from GSE80829 and their classification as per the original publication.

Gene expression data from the CCLE and TCGA were accessed through cBioportal (Cerami et al., 2012; Gao et al., 2013). Moving average trendline of bar plot representing expression level of individual samples was generated using a simple window size of 20. Spearman correlation and the significance was calculated using cor.test function in R.

## Data Availability

The authors declare that all data supporting the findings of this study are available within the article or from the corresponding author upon request. The RNA seq data from the 12 in house melanoma cell lines have been deposited on GEO (accession number: GSE184923 reviewer access token is **yhcxcoicnnsflyl**)

## Ethics Statement

Collection of human adipose tissue was performed according to IRB-HUFA, act number 17/68 granting approval on June ^8t^h, 2017. Adipose tissue samples were collected using protocols approved by the corresponding Ethics Committees and following the legislation. All patients gave written informed consent for the use of their biological samples for research purposes. Fundamental ethical principles and rights promoted by Spain (LOPD 15/1999) and the European Union EU (2000/C364/01) were followed. All patients’ data were processed according to the Declaration of Helsinki (last revision 2013) and Spanish National Biomedical Research Law (14/2007, July 3).

## Acknowledgements

Funding was provided by the Ludwig Institute for Cancer Research (C.R.G, Y.V.G. and P.L.), NIH grants PO1 CA128814-06A1 (Y.V.G., C.R.G.) and R01 CA268597-01 (C.R.G., P.L.), a European Union Marie Curie Training Fellowship (FP7-PEOPLE: PIEF-GA-2013-626098) and European Molecular Biology Organisation (EMBO) Long Term Fellowship (ALTF 800–2013) (A.C-C.), a Ministerio de Economía y Competitividad Grant SAF2016-79837-R and Agencia Estatal de Investigación MCIN/AEI/10.13039/501100011033 (PID2019-104867RB-I00) (C.G.J.) and PID2021-127645OA-I00 (A.C-C); and Comunidad de Madrid PRECICOLON-CM, P2022/BMD-7212 (C.G.J.) and Ayudas Atracción de Talento 2017-T1/BMD-5334 and 2021-5A/BMD-20951 (A.C-C).

## Author Contribution

CG-J. and C.R.G. conceived the project and designed and interpreted experiments. A.C-C., J.M., M.J-O., A.R-S., Y.V-G. and P.L. performed experiments, J.C. and P.L. provided bioinformatic analysis, MC F and MD provided fat explants, C.G-J. and C.R.G. provided resources and supervision. C.G-J. and C.R.G., wrote the manuscript.

## Author information

Correspondence and requests for materials should be sent to CGJ or CRG.

---

**Supplemental Figure 1.**
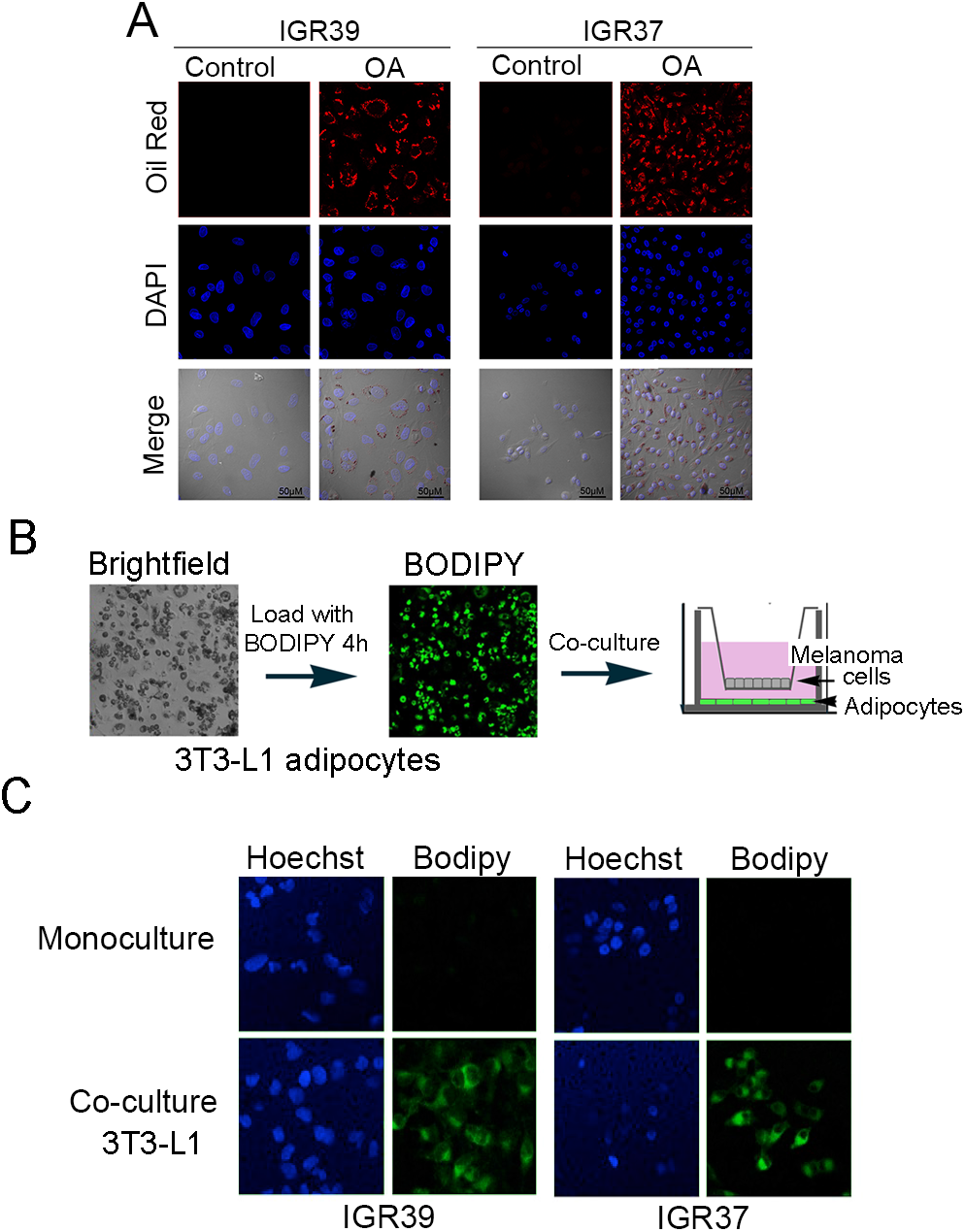
Fatty acid uptake by IGR29 and IGR37 cells. **A.** Oil red O staining of IGR37 and IGR39 melanoma cells after 4 h exposure to oleic acid (OA). Scale bar = 50 µm. **B.** Schematic showing pre-loading of 3T3-L1 adipocytes with BODIPY (green) before removal of free BODIPY by washing and co-culture with melanoma cells in a chamber separated by a membrane that permits passage of small molecules but not cells or cell contact. **C.** BODIPY-FL uptake from pre-loaded 3T3-L1 cells by IGR37 or IGR39 cells co-cultured for 24 h using mono-cultured melanoma cells as a negative control. Both melanoma cell lines are positive, indicating their ability to promote release and uptake from the BODIPY-labelled 3T3-L1 cells.

**Supplemental Figure 2.**
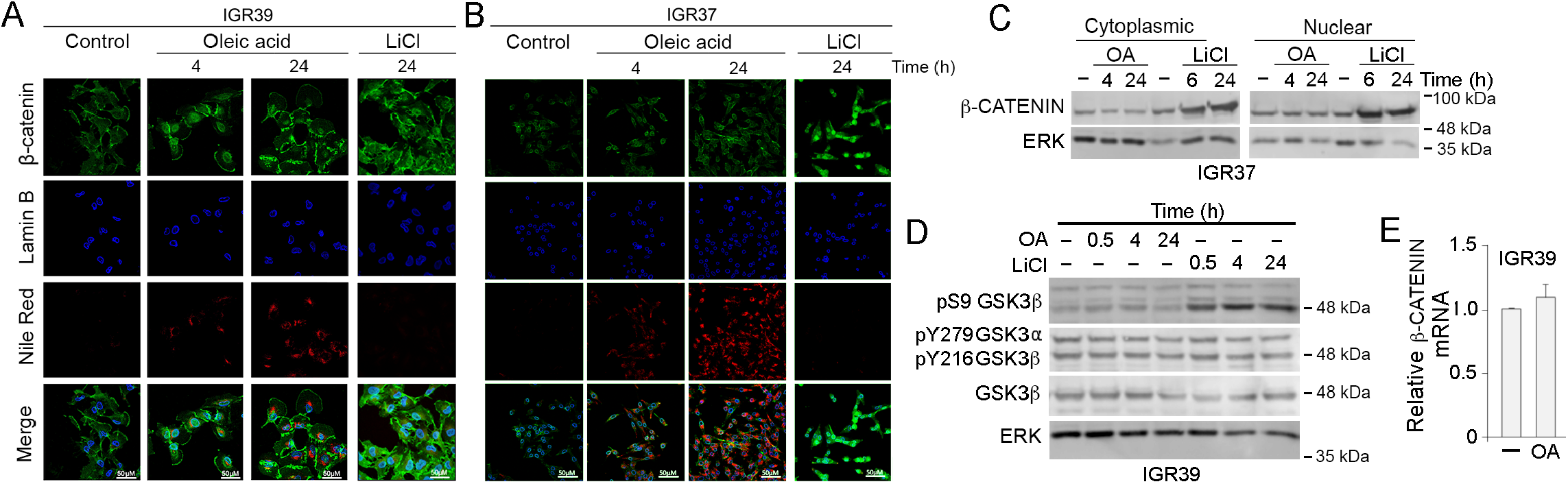
Effect of Oleic acid on β-catenin in IGR37 and IGR39 cells. **A, B.** Immunofluorescence showing subcellular localization of β-catenin following treatment with 100 µM oleic acid or 20 mM LiCl. Anti-Lamin B was used to highlight the nuclear periphery. C. Western blot of fractionated IGR37 cells following treatment with 100 µM oleic acid or 20 mM LiCl. **D**. Western blot over time of IGR39 cells treated with 100 µM oleic acid or 20 mM LiCl. **E.** qRT PCR showing expression of β-catenin mRNA relative to GAPDH in IGR39 cells treated with 100 μM oleic acid for 24 h.

## References

Alicea, G.M., Rebecca, V.W., Goldman, A.R., Fane, M.E., Douglass, S.M., Behera, R., Webster, M.R., Kugel, C.H., 3rd, Ecker, B.L., Caino, M.C., et al. (2020). Changes in Aged Fibroblast Lipid Metabolism Induce Age-Dependent Melanoma Cell Resistance to Targeted Therapy via the Fatty Acid Transporter FATP2. Cancer Discov. 10, 1282–1295. 10.1158/2159-8290.CD-20-0329.

Altea-Manzano, P., Doglioni, G., Liu, Y., Cuadros, A.M., Nolan, E., Fernandez-Garcia, J., Wu, Q., Planque, M., Laue, K.J., Cidre-Aranaz, F., et al. (2023). A palmitate-rich metastatic niche enables metastasis growth via p65 acetylation resulting in pro-metastatic NF-kappaB signaling. Nat Cancer. 10.1038/s43018-023-00513-2.

Anastas, J.N., Kulikauskas, R.M., Tamir, T., Rizos, H., Long, G.V., von Euw, E.M., Yang, P.T., Chen, H.W., Haydu, L., Toroni, R.A., et al. (2014). WNT5A enhances resistance of melanoma cells to targeted BRAF inhibitors. J Clin Invest 124, 2877–2890. 10.1172/JCI70156.

Asiedu, M.K., Beauchamp-Perez, F.D., Ingle, J.N., Behrens, M.D., Radisky, D.C., and Knutson, K.L. (2014). AXL induces epithelial-to-mesenchymal transition and regulates the function of breast cancer stem cells. Oncogene 33, 1316–1324. 10.1038/onc.2013.57.

Aubert, C., Rouge, F., and Galindo, J.R. (1980). Tumorigenicity of human malignant melanocytes in nude mice in relation to their differentiation in vitro. J Natl Cancer Inst 64, 1029–1040.

Auyez, A., Sayan, A.E., Kriajevska, M., and Tulchinsky, E. (2021). AXL Receptor in Cancer Metastasis and Drug Resistance: When Normal Functions Go Askew. Cancers (Basel) 13. 10.3390/cancers13194864.

Bai, J., and Pagano, R.E. (1997). Measurement of spontaneous transfer and transbilayer movement of BODIPY-labeled lipids in lipid vesicles. Biochemistry 36, 8840–8848. 10.1021/bi970145r.

Balaban, S., Shearer, R.F., Lee, L.S., van Geldermalsen, M., Schreuder, M., Shtein, H.C., Cairns, R., Thomas, K.C., Fazakerley, D.J., Grewal, T., et al. (2017). Adipocyte lipolysis links obesity to breast cancer growth: adipocyte-derived fatty acids drive breast cancer cell proliferation and migration. Cancer & Metabolism 5, 1. 10.1186/s40170-016-0163-7.

Bansal, N., Mishra, P.J., Stein, M., DiPaola, R.S., and Bertino, J.R. (2015). Axl receptor tyrosine kinase is up-regulated in metformin resistant prostate cancer cells. Oncotarget 6, 15321–15331. 10.18632/oncotarget.4148.

Boshuizen, J., Koopman, L.A., Krijgsman, O., Shahrabi, A., van den Heuvel, E.G., Ligtenberg, M.A., Vredevoogd, D.W., Kemper, K., Kuilman, T., Song, J.Y., et al. (2018). Cooperative targeting of melanoma heterogeneity with an AXL antibody-drug conjugate and BRAF/MEK inhibitors. Nat Med 24, 203–212. 10.1038/nm.4472.

Cao, X., Tay, A., Guy, G.R., and Tan, Y.H. (1996). Activation and association of Stat3 with Src in v-Src-transformed cell lines. Mol Cell Biol 16, 1595–1603. 10.1128/MCB.16.4.1595.

Carreira, S., Goodall, J., Denat, L., Rodriguez, M., Nuciforo, P., Hoek, K.S., Testori, A., Larue, L., and Goding, C.R. (2006). Mitf regulation of Dia1 controls melanoma proliferation and invasiveness. Genes Dev 20, 3426–3439. 10.1101/gad.406406.

Catalan, V., Gomez-Ambrosi, J., Rodriguez, A., Perez-Hernandez, A.I., Gurbindo, J., Ramirez, B., Mendez-Gimenez, L., Rotellar, F., Valenti, V., Moncada, R., et al. (2014). Activation of noncanonical Wnt signaling through WNT5A in visceral adipose tissue of obese subjects is related to inflammation. J Clin Endocrinol Metab 99, E1407–1417. 10.1210/jc.2014-1191.

Cerami, E., Gao, J., Dogrusoz, U., Gross, B.E., Sumer, S.O., Aksoy, B.A., Jacobsen, A., Byrne, C.J., Heuer, M.L., Larsson, E., et al. (2012). The cBio cancer genomics portal: an open platform for exploring multidimensional cancer genomics data. Cancer Discov. 2, 401–404. 10.1158/2159-8290.CD-12-0095.

Chauhan, J.S., Holzel, M., Lambert, J.P., Buffa, F.M., and Goding, C.R. (2022). The MITF regulatory network in melanoma. Pigment Cell Melanoma Res 35, 517–533. 10.1111/pcmr.13053.

Chocarro-Calvo, A., Garcia-Martinez, J.M., Ardila-Gonzalez, S., De la Vieja, A., and Garcia-Jimenez, C. (2013). Glucose-induced beta-catenin acetylation enhances Wnt signaling in cancer. Mol Cell 49, 474–486. 10.1016/j.molcel.2012.11.022.

Chretien, A., Piront, N., Delaive, E., Demazy, C., Ninane, N., and Toussaint, O. (2008). Increased abundance of cytoplasmic and nuclear caveolin 1 in human diploid fibroblasts in H(2)O(2)-induced premature senescence and interplay with p38alpha(MAPK). FEBS Lett 582, 1685–1692. 10.1016/j.febslet.2008.04.026.

Clevers, H. (2006). Wnt/beta-catenin signaling in development and disease. Cell 127, 469–480.

Colavito, S.A. (2020). AXL as a Target in Breast Cancer Therapy. J Oncol 2020, 5291952. 10.1155/2020/5291952.

Conde-Perez, A., Gros, G., Longvert, C., Pedersen, M., Petit, V., Aktary, Z., Viros, A., Gesbert, F., Delmas, V., Rambow, F., et al. (2015). A caveolin-dependent and PI3K/AKT-independent role of PTEN in β-catenin transcriptional activity. Nat Commun 6, 8093. 10.1038/ncomms9093.

Creedon, H., Byron, A., Main, J., Hayward, L., Klinowska, T., and Brunton, V.G. (2014). Exploring mechanisms of acquired resistance to HER2 (human epidermal growth factor receptor 2)-targeted therapies in breast cancer. Biochem Soc Trans 42, 822–830. 10.1042/BST20140109.

Damsky, W.E., Curley, D.P., Santhanakrishnan, M., Rosenbaum, L.E., Platt, J.T., Gould Rothberg, B.E., Taketo, M.M., Dankort, D., Rimm, D.L., McMahon, M., and Bosenberg, M. (2011). beta-Catenin Signaling Controls Metastasis in Braf-Activated Pten-Deficient Melanomas. Cancer Cell 20, 741–754. S1535-6108(11)00405-3 [pii] 10.1016/j.ccr.2011.10.030.

De Palma, M., Biziato, D., and Petrova, T.V. (2017). Microenvironmental regulation of tumour angiogenesis. Nat Rev Cancer 17, 457–474. 10.1038/nrc.2017.51.

Debruyne, D.N., Bhatnagar, N., Sharma, B., Luther, W., Moore, N.F., Cheung, N.K., Gray, N.S., and George, R.E. (2016). ALK inhibitor resistance in ALK(F1174L)-driven neuroblastoma is associated with AXL activation and induction of EMT. Oncogene 35, 3681–3691. 10.1038/onc.2015.434.

Delmas, V., Beermann, F., Martinozzi, S., Carreira, S., Ackermann, J., Kumasaka, M., Denat, L., Goodall, J., Luciani, F., Viros, A., et al. (2007). Beta-catenin induces immortalization of melanocytes by suppressing p16INK4a expression and cooperates with N-Ras in melanoma development. Genes Dev 21, 2923–2935. 21/22/2923 [pii] 10.1101/gad.450107.

Dirat, B., Bochet, L., Dabek, M., Daviaud, D., Dauvillier, S., Majed, B., Wang, Y.Y., Meulle, A., Salles, B., Le Gonidec, S., et al. (2011). Cancer-associated adipocytes exhibit an activated phenotype and contribute to breast cancer invasion. Cancer Res 71, 2455–2465. 10.1158/0008-5472.CAN-10-3323.

Doege, H., and Stahl, A. (2006). Protein-mediated fatty acid uptake: novel insights from in vivo models. Physiology (Bethesda) 21, 259–268. 10.1152/physiol.00014.2006.

Elkabets, M., Pazarentzos, E., Juric, D., Sheng, Q., Pelossof, R.A., Brook, S., Benzaken, A.O., Rodon, J., Morse, N., Yan, J.J., et al. (2015). AXL mediates resistance to PI3Kα inhibition by activating the EGFR/PKC/mTOR axis in head and neck and esophageal squamous cell carcinomas. Cancer Cell 27, 533–546. 10.1016/j.ccell.2015.03.010.

Falletta, P., Sanchez-del-Campo, L., Chauhan, J., Effern, M., Kenyon, A., Kershaw, C.J., Siddaway, R., Lisle, R., Freter, R., Daniels, M., et al. (2017). Translation reprogramming is an evolutionarily conserved driver of phenotypic plasticity and therapeutic resistance in melanoma. Genes Dev 31, 18–33.

Fogh, J., Fogh, J.M., and Orfeo, T. (1977). One hundred and twenty-seven cultured human tumor cell lines producing tumors in nude mice. J Natl Cancer Inst 59, 221–226.

Gao, C., Chen, G., Kuan, S.F., Zhang, D.H., Schlaepfer, D.D., and Hu, J. (2015). FAK/PYK2 promotes the Wnt/beta-catenin pathway and intestinal tumorigenesis by phosphorylating GSK3beta. eLife 4. 10.7554/eLife.10072.

Gao, J., Aksoy, B.A., Dogrusoz, U., Dresdner, G., Gross, B., Sumer, S.O., Sun, Y., Jacobsen, A., Sinha, R., Larsson, E., et al. (2013). Integrative analysis of complex cancer genomics and clinical profiles using the cBioPortal. Science Signaling 6, pl1. 10.1126/scisignal.2004088.

García-Jiménez, C., and Goding, C.R. (2019). Starvation and Pseudo-Starvation as Drivers of Cancer Metastasis through Translation Reprogramming. Cell Metab 29, 258–267.

García-Jiménez, C., Gutierrez-Salmeron, M., Chocarro-Calvo, A., Garcia-Martinez, J.M., Castano, A., and De la Vieja, A. (2016). From obesity to diabetes and cancer: epidemiological links and role of therapies. Brit J Cancer 114, 716–722. 10.1038/bjc.2016.37.

Gay, C.M., Balaji, K., and Byers, L.A. (2017). Giving AXL the axe: targeting AXL in human malignancy. Brit J Cancer 116, 415–423. 10.1038/bjc.2016.428.

Gerstberger, S., Jiang, Q., and Ganesh, K. (2023). Metastasis. Cell 186, 1564–1579. 10.1016/j.cell.2023.03.003.

Giard, D.J., Aaronson, S.A., Todaro, G.J., Arnstein, P., Kersey, J.H., Dosik, H., and Parks, W.P. (1973). In vitro cultivation of human tumors: establishment of cell lines derived from a series of solid tumors. J Natl Cancer Inst 51, 1417–1423.

Goding, C.R., and Arnheiter, H. (2019). MITF - The first 25 years. Genes Dev 33, 983–1007. doi: 10.1101/gad.324657.119.

Goodall, J., Carreira, S., Denat, L., Kobi, D., Davidson, I., Nuciforo, P., Sturm, R.A., Larue, L., and Goding, C.R. (2008). Brn-2 represses microphthalmia-associated transcription factor expression and marks a distinct subpopulation of microphthalmia-associated transcription factor-negative melanoma cells. Cancer Res 68, 7788–7794. 10.1158/0008-5472.CAN-08-1053.

Hanahan, D. (2022). Hallmarks of Cancer: New Dimensions. Cancer Discov 12, 31–46. 10.1158/2159-8290.CD-21-1059.

Hau, A.M., Gupta, S., Leivo, M.Z., Nakashima, K., Macias, J., Zhou, W., Hodge, A., Wulfkuhle, J., Conkright, B., Bhuvaneshwar, K., et al. (2019). Dynamic Regulation of Caveolin-1 Phosphorylation and Caveolae Formation by Mammalian Target of Rapamycin Complex 2 in Bladder Cancer Cells. Am J Path 189, 1846–1862. 10.1016/j.ajpath.2019.05.010.

Henderson, G.C. (2021). Plasma Free Fatty Acid Concentration as a Modifiable Risk Factor for Metabolic Disease. Nutrients 13. 10.3390/nu13082590.

Herlyn, M., Thurin, J., Balaban, G., Bennicelli, J.L., Herlyn, D., Elder, D.E., Bondi, E., Guerry, D., Nowell, P., Clark, W.H., and, et al. (1985). Characteristics of cultured human melanocytes isolated from different stages of tumor progression. Cancer Res 45, 5670–5676.

Hirata, E., Girotti, M.R., Viros, A., Hooper, S., Spencer-Dene, B., Matsuda, M., Larkin, J., Marais, R., and Sahai, E. (2015). Intravital Imaging Reveals How BRAF Inhibition Generates Drug-Tolerant Microenvironments with High Integrin beta1/FAK Signaling. Cancer Cell 27, 574–588. 10.1016/j.ccell.2015.03.008.

Hodson, L., Skeaff, C.M., and Fielding, B.A. (2008). Fatty acid composition of adipose tissue and blood in humans and its use as a biomarker of dietary intake. Prog Lipid Res 47, 348–380. 10.1016/j.plipres.2008.03.003.

Hoek, K., and Goding, C.R. (2010). Cancer stem cells versus phenotype switching in melanoma. Pigment Cell Melanoma Res. 23, 746–759.

Hollander, D.M., Devereux, D.F., Taylor, C.G., and Taylor, D.D. (1986). Demonstration of lipolytic activity from cultured human melanoma cells. J Surg Res 40, 445–449.

Hoy, A.J., Nagarajan, S.R., and Butler, L.M. (2021). Tumour fatty acid metabolism in the context of therapy resistance and obesity. Nat Rev Cancer 21, 753–766. 10.1038/s41568-021-00388-4.

Hugo, W., Zaretsky, J.M., Sun, L., Song, C., Moreno, B.H., Hu-Lieskovan, S., Berent-Maoz, B., Pang, J., Chmielowski, B., Cherry, G., et al. (2016). Genomic and Transcriptomic Features of Response to Anti-PD-1 Therapy in Metastatic Melanoma. Cell 165, 35–44. 10.1016/j.cell.2016.02.065.

Insull, W., Jr., and Bartsch, G.E. (1967). Fatty acid composition of human adipose tissue related to age, sex, and race. Am J Clin Nutr 20, 13–23. 10.1093/ajcn/20.1.13.

Kalkavan, H., Chen, M.J., Crawford, J.C., Quarato, G., Fitzgerald, P., Tait, S.W.G., Goding, C.R., and Green, D.R. (2022). Sublethal cytochrome c release generates drug-tolerant persister cells. Cell 185, 3356–3374 e3322. 10.1016/j.cell.2022.07.025.

Kmiecik, T.E., Johnson, P.J., and Shalloway, D. (1988). Regulation by the autophosphorylation site in overexpressed pp60c-src. Mol Cell Biol 8, 4541–4546. 10.1128/mcb.8.10.4541.

Kokatnur, M.G., Oalmann, M.C., Johnson, W.D., Malcom, G.T., and Strong, J.P. (1979). Fatty acid composition of human adipose tissue from two anatomical sites in a biracial community. Am J Clin Nutr 32, 2198–2205. 10.1093/ajcn/32.11.2198.

Konieczkowski, D.J., Johannessen, C.M., Abudayyeh, O., Kim, J.W., Cooper, Z.A., Piris, A., Frederick, D.T., Barzily-Rokni, M., Straussman, R., Haq, R., et al. (2014). A melanoma cell state distinction influences sensitivity to MAPK pathway inhibitors. Cancer Discov 4, 816–827. 10.1158/2159-8290.CD-13-0424.

Kwan, H.Y., Fu, X., Liu, B., Chao, X., Chan, C.L., Cao, H., Su, T., Tse, A.K., Fong, W.F., and Yu, Z.-L.L. (2014). Subcutaneous adipocytes promote melanoma cell growth by activating the Akt signaling pathway: role of palmitic acid. J Biol Chem 289, 30525–30537. 10.1074/jbc.M114.593210.

Lazar, I., Clement, E., Dauvillier, S., Milhas, D., Ducoux-Petit, M., LeGonidec, S., Moro, C., Soldan, V., Dalle, S., Balor, S., et al. (2016). Adipocyte Exosomes Promote Melanoma Aggressiveness through Fatty Acid Oxidation: A Novel Mechanism Linking Obesity and Cancer. Cancer Res 76, 4051–4057. 10.1158/0008-5472.CAN-16-0651.

Lee, H.J., Chen, Z., Collard, M., Chen, F., J.G., C., Wu, M., Alani, R.M., and Chen, J.-X. (2021). Multimodal metabolic imaging reveals pigment reduction and lipid accumulation in metastatic melanoma. BMEF 2021, 986Q123.

Li, S., Seitz, R., and Lisanti, M.P. (1996). Phosphorylation of caveolin by src tyrosine kinases. The alpha-isoform of caveolin is selectively phosphorylated by v-Src in vivo. J Biol Chem 271, 3863–3868.

Livak, K.J., and Schmittgen, T.D. (2001). Analysis of relative gene expression data using real-time quantitative PCR and the 2(-Delta Delta C(T)) Method. Methods 25, 402–408. 10.1006/meth.2001.1262.

Louphrasitthiphol, P., Chauhan, J., and Goding, C.R. (2020). ABCB5 is activated by MITF and beta-catenin and is associated with melanoma differentiation. Pigment Cell Melanoma Res 33, 112–118. 10.1111/pcmr.12830.

Louphrasitthiphol, P., Ledaki, I., Chauhan, J., Falletta, P., Siddaway, R., Buffa, F.M., Mole, D.R., Soga, T., and Goding, C.R. (2019). MITF controls the TCA cycle to modulate the melanoma hypoxia response. Pigment Cell Melanoma Res 32, 792–808. 10.1111/pcmr.12802.

Lumaquin-Yin, D., Montal, E., Johns, E., Baggiolini, A., Huang, T.H., Ma, Y., LaPlante, C., Suresh, S., Studer, L., and White, R.M. (2023). Lipid droplets are a metabolic vulnerability in melanoma. Nat Commun 14, 3192. 10.1038/s41467-023-38831-9.

Lyssiotis, C.A., and Kimmelman, A.C. (2017). Metabolic Interactions in the Tumor Microenvironment. Trends Cell Biol 27, 863–875. 10.1016/j.tcb.2017.06.003.

Martinez-Outschoorn, U.E., Lisanti, M.P., and Sotgia, F. (2014). Catabolic cancer-associated fibroblasts transfer energy and biomass to anabolic cancer cells, fueling tumor growth. Seminars in Cancer Biol 25, 47–60. 10.1016/j.semcancer.2014.01.005.

Mattern, H.M., Raikar, L.S., and Hardin, C.D. (2009). The effect of caveolin-1 (Cav-1) on fatty acid uptake and CD36 localization and lipotoxicity in vascular smooth muscle (VSM) cells. Int J Physiol Pathophysiol Pharmacol 1, 1–14.

Morfoisse, F., De Toni, F., Nigri, J., Hosseini, M., Zamora, A., Tatin, F., Pujol, F., Sarry, J.E., Langin, D., Lacazette, E., et al. (2021). Coordinating Effect of VEGFC and Oleic Acid Participates to Tumor Lymphangiogenesis. Cancers (Basel) 13. 10.3390/cancers13122851.

Muller, C., Nieto, L., and Valet, P. (2013). Unraveling the Local Influence of Tumor-Surrounding Adipose Tissue on Tumor Progression: Cellular and Molecular Actors Involved. In Adipose Tissue and Cancer, M.G. Kolonin, ed.

Muller, J., Krijgsman, O., Tsoi, J., Robert, L., Hugo, W., Song, C., Kong, X., Possik, P.A., Cornelissen-Steijger, P.D., Foppen, M.H., et al. (2014). Low MITF/AXL ratio predicts early resistance to multiple targeted drugs in melanoma. Nat Commun 5, 5712. 10.1038/ncomms6712.

Nakajima, E.C., and Van Houten, B. (2013). Metabolic symbiosis in cancer: refocusing the Warburg lens. Mol Carcinogenesis 52, 329–337. 10.1002/mc.21863.

Nieman, K.M., Kenny, H.A., Penicka, C.V., Ladanyi, A., Buell-Gutbrod, R., Zillhardt, M.R., Romero, I.L., Carey, M.S., Mills, G.B., Hotamisligil, G.S., et al. (2011). Adipocytes promote ovarian cancer metastasis and provide energy for rapid tumor growth. Nat Med 17, 1498–1503. nm.2492 [pii] 10.1038/nm.2492.

Ortiz, R., Diaz, J., Diaz, N., Lobos-Gonzalez, L., Cardenas, A., Contreras, P., Diaz, M.I., Otte, E., Cooper-White, J., Torres, V., et al. (2016). Extracellular matrix-specific Caveolin-1 phosphorylation on tyrosine 14 is linked to augmented melanoma metastasis but not tumorigenesis. Oncotarget 7, 40571–40593. 10.18632/oncotarget.9738.

Pandey, V., Vijayakumar, M.V., Ajay, A.K., Malvi, P., and Bhat, M.K. (2012). Diet-induced obesity increases melanoma progression: involvement of Cav-1 and FASN. Int. J Cancer 130, 497–508. 10.1002/ijc.26048.

Pascual, G., Avgustinova, A., Mejetta, S., Martín, M., Castellanos, A., Attolini, C.S.-O., Berenguer, A., Prats, N., Toll, A., Hueto, J.A., et al. (2016). Targeting metastasis-initiating cells through the fatty acid receptor CD36. Nature 541, 41–45. doi:10.1038/nature20791.

Pavlova, N.N., and Thompson, C.B. (2016). The Emerging Hallmarks of Cancer Metabolism. Cell Metab 23, 27–47. 10.1016/j.cmet.2015.12.006.

Pohl, J., Ring, A., Ehehalt, R., Herrmann, T., and Stremmel, W. (2004). New concepts of cellular fatty acid uptake: role of fatty acid transport proteins and of caveolae. Proc Nutr Soc 63, 259–262. 10.1079/PNS2004341.

Pohl, J., Ring, A., Korkmaz, U., Ehehalt, R., and Stremmel, W. (2005). FAT/CD36-mediated long-chain fatty acid uptake in adipocytes requires plasma membrane rafts. Mol Biol Cell 16, 24–31. 10.1091/mbc.e04-07-0616.

Rabbani, P., Takeo, M., Chou, W., Myung, P., Bosenberg, M., Chin, L., Taketo, M.M., and Ito, M. (2011). Coordinated activation of Wnt in epithelial and melanocyte stem cells initiates pigmented hair regeneration. Cell 145, 941–955. S0092-8674(11)00530-7 [pii] 10.1016/j.cell.2011.05.004.

Rambow, F., Marine, J.C., and Goding, C.R. (2019). Melanoma plasticity and phenotypic diversity: therapeutic barriers and opportunities Genes Dev 33, 1295–1318.

Rambow, F., Rogiers, A., Marin-Bejar, O., Aibar, S., Femel, J., Dewaela, M., Karras, P., Brown, D., Chang, Y.H., Debiec-Rychter, M., et al. (2018). Towards minimal residual disease-directed therapy in melanoma. Cell 174, 843–855.

Riesenberg, S., Groetchen, A., Siddaway, R., Bald, T., Reinhardt, J., Smorra, D., Kohlmeyer, J., Renn, M., Phung, B., Aymans, P., et al. (2015). MITF and c-Jun antagonism interconnects melanoma dedifferentiation with pro-inflammatory cytokine responsiveness and myeloid cell recruitment. Nat Commun 6, 8755 DOI:8710.1038 ncomss9755. 10.1038/ncomms9755.

Ring, A., Le Lay, S., Pohl, J., Verkade, P., and Stremmel, W. (2006). Caveolin-1 is required for fatty acid translocase (FAT/CD36) localization and function at the plasma membrane of mouse embryonic fibroblasts. Biochimica Biophysica Acta 1761, 416–423. 10.1016/j.bbalip.2006.03.016.

Sandoval, A., Chokshi, A., Jesch, E.D., Black, P.N., and Dirusso, C.C. (2010). Identification and characterization of small compound inhibitors of human FATP2. Biochem Pharmacol 79, 990–999. 10.1016/j.bcp.2009.11.008.

Sanna, E., Miotti, S., Mazzi, M., De Santis, G., Canevari, S., and Tomassetti, A. (2007). Binding of nuclear caveolin-1 to promoter elements of growth-associated genes in ovarian carcinoma cells. Exp Cell Res 313, 1307–1317. 10.1016/j.yexcr.2007.02.005.

Shao, H., Teramae, D., and Wells, A. (2023). Axl contributes to efficient migration and invasion of melanoma cells. PloS one 18, e0283749. 10.1371/journal.pone.0283749.

Sharma, S.V., Lee, D.Y., Li, B., Quinlan, M.P., Takahashi, F., Maheswaran, S., McDermott, U., Azizian, N., Zou, L., Fischbach, M.A., et al. (2010). A chromatin-mediated reversible drug-tolerant state in cancer cell subpopulations. Cell 141, 69–80. S0092-8674(10)00180-7 [pii] 10.1016/j.cell.2010.02.027.

Smolle, J., Woltsche, I., Hofmann-Wellenhof, R., Haas, J., and Kerl, H. (1995). Pathology of tumor-stroma interaction in melanoma metastatic to the skin. Hum Pathol 26, 856–861.

Sommer, L. (2011). Generation of Melanocytes from Neural Crest Cells. Pigment Cell Melanoma Res. 10.1111/j.1755-148X.2011.00834.x.

Sonveaux, P., Vegran, F., Schroeder, T., Wergin, M.C., Verrax, J., Rabbani, Z.N., De Saedeleer, C.J., Kennedy, K.M., Diepart, C., Jordan, B.F., et al. (2008). Targeting lactate-fueled respiration selectively kills hypoxic tumor cells in mice. J Clin Invest 118, 3930–3942. 10.1172/JCI36843.

Sousa, C.M., Biancur, D.E., Wang, X., Halbrook, C.J., Sherman, M.H., Zhang, L., Kremer, D., Hwang, R.F., Witkiewicz, A.K., Ying, H., et al. (2016). Pancreatic stellate cells support tumour metabolism through autophagic alanine secretion. Nature 536, 479–483. 10.1038/nature19084.

Tagawa, A., Mezzacasa, A., Hayer, A., Longatti, A., Pelkmans, L., and Helenius, A. (2005). Assembly and trafficking of caveolar domains in the cell: caveolae as stable, cargo-triggered, vesicular transporters. J Cell Biol 170, 769–779. 10.1083/jcb.200506103.

Tsoi, J., Robert, L., Paraiso, K., Galvan, C., Sheu, K.M., Lay, J., Wong, D.J.L., Atefi, M., Shirazi, R., Wang, X., et al. (2018). Multi-stage Differentiation Defines Melanoma Subtypes with Differential Vulnerability to Drug-Induced Iron-Dependent Oxidative Stress. Cancer Cell 33, 890–904 e895. 10.1016/j.ccell.2018.03.017.

Ubellacker, J.M., Tasdogan, A., Ramesh, V., Shen, B., Mitchell, E.C., Martin-Sandoval, M.S., Gu, Z., McCormick, M.L., Durham, A.B., Spitz, D.R., et al. (2020). Lymph protects metastasizing melanoma cells from ferroptosis. Nature 585, 113–118. 10.1038/s41586-020-2623-z.

Veeman, M.T., Slusarski, D.C., Kaykas, A., Louie, S.H., and Moon, R.T. (2003). Zebrafish prickle, a modulator of noncanonical Wnt/Fz signaling, regulates gastrulation movements. Curr Biol 13, 680–685. 10.1016/s0960-9822(03)00240-9.

Vivas-Garcia, Y., Falletta, P., Liebing, J., Louphrasitthiphol, P., Feng, Y., Chauhan, J., Scott, D.A., Glodde, N., Chocarro-Calvo, A., Bonham, S., et al. (2020). Lineage-Restricted Regulation of SCD and Fatty Acid Saturation by MITF Controls Melanoma Phenotypic Plasticity. Mol Cell 77, 120–139. 10.1016/j.molcel.2019.10.014.

Wagner, M., Bjerkvig, R., Wiig, H., Melero-Martin, J.M., Lin, R.Z., Klagsbrun, M., and Dudley, A.C. (2012). Inflamed tumor-associated adipose tissue is a depot for macrophages that stimulate tumor growth and angiogenesis. Angiogenesis 15, 481–495. 10.1007/s10456-012-9276-y.

Wang, C., Jin, H., Wang, N., Fan, S., Wang, Y., Zhang, Y., Wei, L., Tao, X., Gu, D., Zhao, F., et al. (2016). Gas6/Axl Axis Contributes to Chemoresistance and Metastasis in Breast Cancer through Akt/GSK-3beta/beta-catenin Signaling. Theranostics 6, 1205–1219. 10.7150/thno.15083.

Weeraratna, A.T., Jiang, Y., Hostetter, G., Rosenblatt, K., Duray, P., Bittner, M., and Trent, J.M. (2002). Wnt5a signaling directly affects cell motility and invasion of metastatic melanoma. Cancer Cell 1, 279–288. 10.1016/s1535-6108(02)00045-4.

Widlund, H.R., Horstmann, M.A., Price, E.R., Cui, J., Lessnick, S.L., Wu, M., He, X., and Fisher, D.E. (2002). Beta-catenin-induced melanoma growth requires the downstream target Microphthalmia-associated transcription factor. J Cell Biol 158, 1079–1087. 10.1083/jcb.200202049.

Yang, W., Xia, Y., Ji, H., Zheng, Y., Liang, J., Huang, W., Gao, X., Aldape, K., and Lu, Z. (2011). Nuclear PKM2 regulates beta-catenin transactivation upon EGFR activation. Nature 480, 118–122. 10.1038/nature10598.

Zakut, R., Perlis, R., Eliyahu, S., Yarden, Y., Givol, D., Lyman, S.D., and Halaban, R. (1993). KIT ligand (mast cell growth factor) inhibits the growth of KIT-expressing melanoma cells. Oncogene 8, 2221–2229.

Zhang, M., Di Martino, J.S., Bowman, R.L., Campbell, N.R., Baksh, S.C., Simon-Vermot, T., Kim, I.S., Haldeman, P., Mondal, C., Yong-Gonzales, V., et al. (2018). Adipocyte-Derived Lipids Mediate Melanoma Progression via FATP Proteins. Cancer Discov 8, 1006–1025. 10.1158/2159-8290.CD-17-1371.

Zhang, Z., Lee, J.C., Lin, L., Olivas, V., Au, V., LaFramboise, T., Abdel-Rahman, M., Wang, X., Levine, A.D., Rho, J.K., et al. (2012). Activation of the AXL kinase causes resistance to EGFR-targeted therapy in lung cancer. Nat Genet 44, 852–860. 10.1038/ng.2330.

Zhou, L., Liu, X.D., Sun, M., Zhang, X., German, P., Bai, S., Ding, Z., Tannir, N., Wood, C.G., Matin, S.F., et al. (2016). Targeting MET and AXL overcomes resistance to sunitinib therapy in renal cell carcinoma. Oncogene 35, 2687–2697. 10.1038/onc.2015.343.

Zoico, E., Darra, E., Rizzatti, V., Budui, S., Franceschetti, G., Mazzali, G., Rossi, A.P., Fantin, F., Menegazzi, M., Cinti, S., and Zamboni, M. (2016). Adipocytes WNT5a mediated dedifferentiation: a possible target in pancreatic cancer microenvironment. Oncotarget 7, 20223–20235. 10.18632/oncotarget.7936.

